# Adult spiny mice (*Acomys*) exhibit endogenous cardiac recovery in response to myocardial infarction

**DOI:** 10.1101/2020.09.29.317388

**Authors:** Hsuan Peng, Kazuhiro Shindo, Renée R. Donahue, Erhe Gao, Brooke M. Ahern, Bryana M. Levitan, Himi Tripathi, David Powell, Ahmed Noor, Jonathan Satin, Ashley W. Seifert, Ahmed Abdel-Latif

**Affiliations:** Saha Cardiovascular Research Center, College of Medicine, University of Kentucky; The Center for Translational Medicine, Lewis Katz School of Medicine, Temple University, Philadelphia, PA, USA; Department of Physiology, College of Medicine, University of Kentucky; Gill Heart and Vascular Institute and Division of Cardiovascular Medicine, University of Kentucky; Department of Biology, University of Kentucky; The Lexington VA Medical Center

**Author notes:** To whom correspondence should be addressed: Jonathan Satin; Ashley W. Seifert and Ahmed Abdel-Latif. These authors contributed equally to the study (Hsuan Peng, Kazuhiro Shindo).

## Abstract

Complex tissue regeneration is extremely rare among adult mammals. An exception, however, is the superior tissue healing of multiple organs in spiny mice (*Acomys*). While *Acomys* species exhibit the remarkable ability to heal complex tissue with minimal scarring, little is known about their cardiac structure and response to cardiac injury. In this study, we first examined baseline *Acomys* cardiac anatomy and function in comparison with the commonly used laboratory *Mus* strains (C57BL6 and SWR). Our results demonstrated comparable cardiac anatomy and function between *Acomys* and *Mus*, but *Acomys* exhibited a higher percentage of cardiomyocytes exhibiting immature characteristics. In response to myocardial infarction, all animals experienced a comparable level of initial cardiac damage. However, *Acomys* demonstrated superior ischemic tolerance and cytoprotection in response to injury as evidenced by cardiac functional stabilization, higher survival rate and smaller scar size 50 days after injury compared to the inbred and outbred mouse strains. Overall, these findings demonstrate augmented myocardial preservation in spiny mice post-MI and establish *Acomys* as a new adult mammalian model for cardiac research.

## Introduction

Significant clinical advances for heart revascularization and next generation medical therapies have improved the mortality rate after myocardial infarction (MI).^1^ The initial ischemic insult, even with timely revascularization, is followed by microvascular damage and infarct expansion leading to exacerbated damage.^2^ Unfortunately, no therapies exist to limit myocardial damage from MI and the ensuing infarct expansion where millions of recovering patients progress to develop heart failure (HF). This is because the normal healing response to tissue injury in adult mammals is fibrotic repair; an effective short-term strategy, but one that leads to compromised tissue function. Fibrotic repair is the default strategy in the adult heart where it is the leading cause of the clinical HF epidemic.^3^ Multiple therapies developed for limiting infarct expansion or cardiac repair based on animal models have achieved only modest clinical success in humans. A failure to translate basic research findings into successful clinical treatments can be traced, in part, to a lack of adult mammalian models possessing the ability of myocardial preservation.

In contrast to humans^4-6^, the ability to regenerate injured organs is widespread among vertebrates. Fishes, newts, and salamanders have extensive regenerative ability, and can functionally replace heart tissue after amputation or severe injury by expediting revascularization^7^ and by mobilizing a highly proliferative cardiomyocyte pool^8,9^. Specifically, zebrafish are a well-characterized model for adult cardiac regeneration with the documented ability to recover from a plethora of heart injuries including apical resection,^10^ cryo-injury,^11^ and coronary artery ligation. Neonatal mice are also capable of limiting the initial damage and repairing the myocardium after excision and ischemic injury during the first few days after birth.^12^ In contrast, most adult mammals generally exhibit poor regenerative capacity, especially as it pertains to recovering from heart damage. Spiny mice (*Acomys spp*.) are murid rodents found throughout Africa, the Middle East and Western Asia. These rodents exhibit a number of special traits;^13^ tantamount among them is the ability to regenerate skin, complex tissue^14-19^ and nephric tissue.^20^ At present, how spiny mice respond to heart injury and the extent to which they can recover remains unknown, although a recent review on the topic raised the intriguing possibility that they might exhibit enhanced cardiac healing.^21^

To rigorously examine how spiny mice respond to heart damage, we first detail *Acomys* heart structure and cardiac function in comparison to the most widely used inbred laboratory *Mus* model of cardiac injury (C57BL6). For comparative purposes we also used the Swiss Webster outbred strain (SWR) to account for greater genetic diversity within our spiny mouse population. This cross-species characterization indicates similar cardiac structure and comparable baseline function among spiny mice and the examined mouse strains. We next conducted a comparative analysis of these animals in response to permanent coronary artery ligation (MI). In contrast to the mouse strains studied, *Acomys* showed significantly improved survival and enhanced myocardial preservation following ischemic injury. The ability of *Acomys* to recover from ischemic injury is coincident with greatly improved angiogenesis rarely seen in adult mammals and unique cardiomyocyte characteristics (higher percentage of small, mono-nucleated and diploid CMs, T-type calcium channels). These findings in *Acomys* support a rapid angiogenic response into fibrotic tissue which limits scar spread and enhances ischemic tolerance in response to MI.

## Methods

### Detailed methods are described in the Supplementary Methods

#### Animal care

Male *Acomys cahirinus* (6-8 months old, sexually mature animals) were obtained from our in-house breeding colony. Male *Mus musculus* (8-12 weeks, sexually mature animals) C57BL6 were obtained from the Jackson Laboratory, Bar Harbor, ME and outbred SWR were obtained from Charles River. Both strains were housed at the University of Kentucky, Lexington, KY. All animal procedures were approved by the University of Kentucky Institutional Animal Care and Use Committee (IACUC #: 2019-3254, 2013-1155 and 2011-0889).

#### Murine model of myocardial infarction

MI surgery was performed using a minimally invasive permanent left anterior descending artery (LAD) ligation model as previously described.^22^

#### Cardiomyocyte isolation and measurements

Ventricular cardiomyocytes were isolated using the Langendorff method as previously described^23^ and are further detailed in the supplementary methods. Images used to analyze cardiomyocyte surface area and nucleation were acquired using an Olympus IX-71 and Olympus BX53 microscopes (Olympus, Tokyo, Japan) at 10x magnification. Olympus cellSens software was used to determine cell surface area and number of nuclei. Nuclear volume and ploidy were determined by Imaris reconstruction of z-stack images acquired using a Nikon Ti, A1 confocal microscope (Nikon, Japan) with a step size of 3 μm and 40x oil immersion objective.

#### Echocardiography

Echocardiography was acquired using a Vevo 3100 (VisualSonics, Toronto, Canada) equipped with a MX550D 25-55 MHz linear array transducer. Cardiac function was assessed as previously described.^22^ Echocardiographic imaging and analyses were performed by a blinded investigator.

#### Cardiac Magnetic Resonance Imaging

Cardiac magnetic resonance imaging (CMR) was performed on a 7-Tesla ClinScan system (Bruker, Ettlingen, Germany, http://www.bruker.com) equipped with a 4-element phased-array cardiac coil and a gradient system with a maximum strength of 450 mT/m and a maximum slew rate of 4,500 mT/m/s. Studies were performed within 72 hours of MI and at 14 days following injury. For late gadolinium-enhanced magnetic resonance imaging, a 0.6 mmol/kg bolus of gadolinium-diethylene triamine pentaacetic acid (Gd-DTPA; Gadavist, Bayer Health Care, Whippany, NJ) was injected using the intraperitoneal route.

#### Electrophysiological recordings and calcium transients

I_Ca,L_ was recorded from freshly isolated cardiomyocytes using the whole-cell configuration of the patch clamp technique, and cytosolic Ca^2+^ transients were collected using fura2-AM as previously described.^23^

#### Tissue collection and Processing

Tissue from all species was collected and weighed immediately at sacrifice. The dry lung weight was collected after incubation at 65° C for three days. Hearts were perfused with PBS (VWR International) followed by 4% PFA (VWR International) fixation via cannulation of the ascending aorta. Hearts were post-fixed overnight at 4^°^C. Hearts were then cut in half along the long axis at the level of the ligation and transferred to 70% ethanol until embedding. Tissues were then paraffin embedded and sectioned at 5um for immunofluorescence and at 8um for Masson’s Trichrome and Picrosirius Red staining.

#### Histology

Paraffin-embedded heart sections were stained with Masson’s Trichrome or Picrosirius Red. Morphometric analyses and image acquisition were performed using a Olympus BX53 microscope (Olympus, Tokyo, Japan) and NIH ImageJ 1.46R software. Quantification of percent fibrosis and scar thickness are detailed in the supplementary methods. Scar collagen fiber quality was examined under 40x magnification. Cellular density in scar tissue was determined by quantifying DAPI+ nuclei in the infarct region at 60x magnification. Infarct size was also quantified three days post-MI from Masson’s Trichrome stained slides as the percentage of left ventricle affected. These images were acquired on Zeiss Axioscan slide scanner at 10x magnification. Images were quantified using Nis-Elements software (Nikon).

#### Immunofluorescence

Immunofluorescence assessments were carried out on deparaffinized and rehydrated sections as previously described.^22^ Images were taken with a 40x Oil Immersion lens on a Nikon A1 Confocal Microscope in the University of Kentucky Confocal Microscopy facility. Isolectin staining was performed on tissues from day 50 post-MI and was quantified at both the peri-infarct border and the center of the scar and is presented as total capillary density per mm^2^. Baseline isolectin measurements were taken at comparable location to the peri-infarct area. A complete list of antibodies is included in the supplemental data.

#### Statistics

Values are expressed as mean ± standard error of mean (SEM). Data were analyzed using a Student’s t-test, one-way ANOVA, repeated-measures ANOVA or mixed-effects ANOVA with Tukey correction to compare data across species as appropriate. Animal numbers are presented as a range based on the number of animals included in each analysis. These numbers are included in the figure legends. All statistical analyses were performed using the Prism 9 software package (GraphPad, La Jolla, CA).

## Results

### Heart and coronary tree anatomy are similar among *Acomys* and *Mus* (C57BL6 and SWR)

To assess whether interpretable comparisons could be made across species, we first characterized hearts from *Acomys* and the most commonly used laboratory mouse (*Mus*) strain: C57BL6. To control for lifespan differences between species, we used 6 month old, sexually mature *Acomys* and 8-12 week old *Mus* for these studies. In an attempt to control for increased genetic diversity in *Acomys* as well as their larger size, we also included an outbred Swiss Webster strain (SWR – Charles River). As these outbred mice have not been evaluated for myocardial ischemia research, this strain also allowed us to assess whether they shared similarities with the SWR/J strain that is known to exhibit a high percentage of mononuclear cardiomyocytes (CMs) that contribute to enhanced cardiac recovery after ischemic injury.^24^ First, we set out to determine whether baseline differences in cardiac gravimetrics exist between *Acomys* and *Mus* (C57BL6 and SWR). We assessed heart weight (HW) and normalized it to body size (body weight [BW] and tibia length [TL]) to account for overall size differences between species (Fig. 1A-D, and Supplementary Table 1). Although *Acomys* demonstrated a higher mean HW compared to C57BL6 (Fig. 1B), normalized to BW, *Acomys* was not significantly different compared to SWR and exhibited only a slighly lower HW/BW compared to C57BL6 (Fig. 1C). SWR mice exhibited a slightly higher HW compared to *Acomys* and C57BL6 when HW was normalized to TL (Fig. 1D). *Acomys* demonstrated heavier wet lung weight compared to *Mus* (C57BL6 and SWR) (Supplementary Fig. 1A), but lower wet and dry lung weight after normalization by BW (Supplementary Fig. 1B, E). Importantly, dry lung weight before normalization by TL and dry and wet lung weight after normalization by TL were not significantly different across species (Supplementary Fig. 1D-F). The complete data is summarized in Supplementary Table 1.

**Figure 1.**
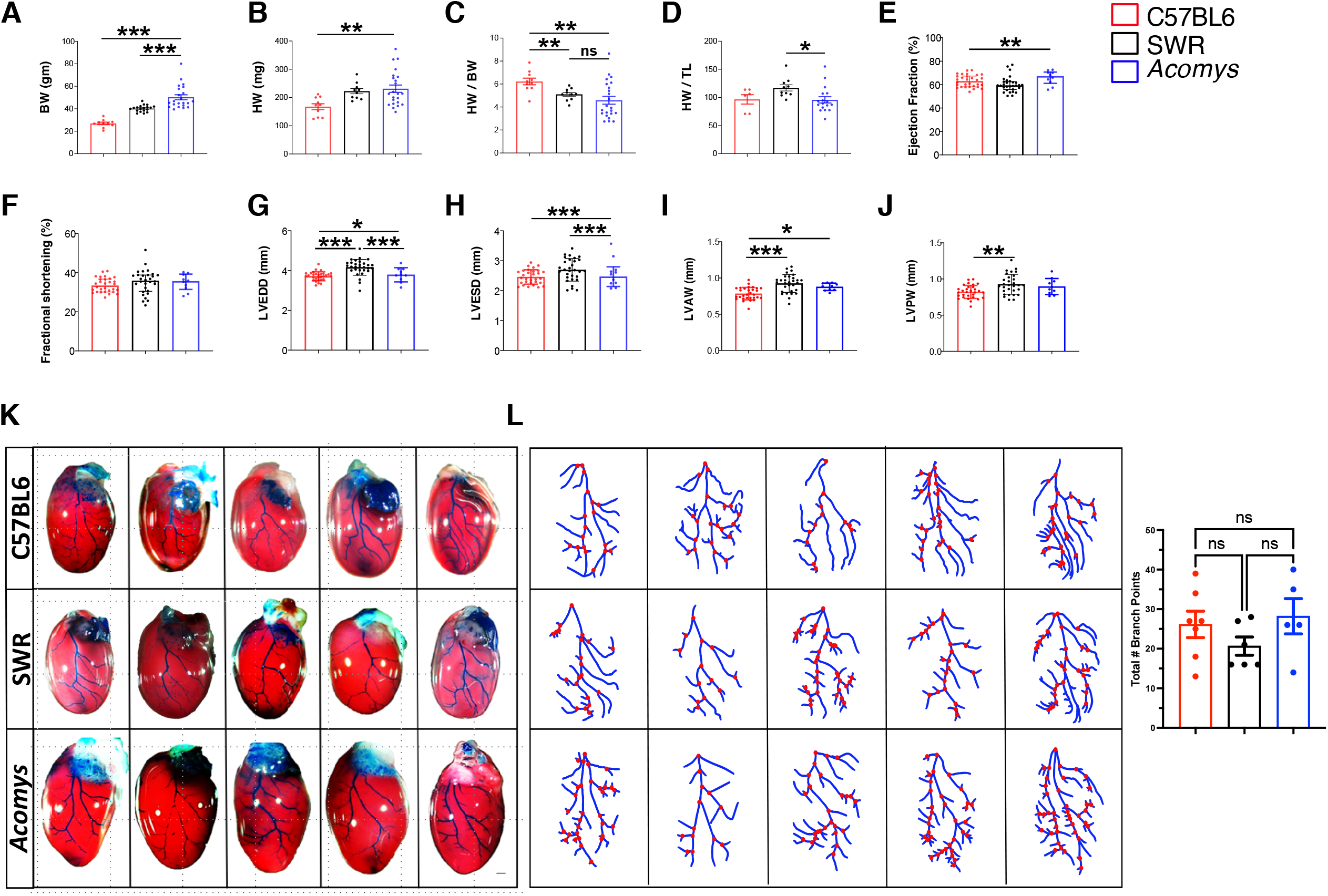
Characteristics of *Acomys* heart physiology and anatomy at baseline. (A-J) Measures of body weight (BW) and heart weight in *Mus*-C57BL6 (N = 10), *Mus*-SWR (N = 19) and *Acomys* (N = 22). Analyses demonstrate comparable heart weight across species when normalized by body weight (BW) and tibia length (TL) (values are means ± S.E.M, \**P* < 0.05, \*\**P*<0.01, and \*\*\**P*<0.001 by one-way ANOVA and Dunnett correction with *Acomys* as control). Ejection fraction (F=4.1, \**P*<0.05) (E), fractional shortening (F=0.96, *P*>0.05) (F), left ventricular end-diastolic diameter (LVEDD) (F=14.8, \*\**P*<0.01) (G), left ventricular end-systolic diameter (LVESD) (F=4.6) (H), left ventricular anterior wall (LVAW) (F=14.9, \**P*<0.05) (I), left ventricular posterior wall (LVPW) (F=6.4) (J) (N = 30 C57BL6-*Mus*, 10 SWR-*Mus* and 10 *Acomys*). (K) Representative images showing comparable left anterior descending artery coronary anatomy across species using intra-aortic infusion of Batson’s 17 polymer mixture and corrosion casting (N = 5 each group, scale bar =1 mm). (L) Quantification of coronary branching points of the arterial supply of the left ventricle across species. This analysis shows comparable number of branching points across species with numerically higher (but not statistically significant) branching points in C57BL6-*Mus* and *Acomys* compared to SWR-*Mus*. Detailed statistical output for Figure 1 can be found in the Supplementary Information.

Next, we compared cardiac functional parameters between the three strains and observed comparable baseline cardiac function under physiological conditions across species. Importantly, the global cardiac function as presented by left ventricular ejection fraction and fractional shortening were similar between all groups (Fig. 1E-F). Measures of left ventricular cavity diameter were also comparable between species except for SWR-*Mus* showing slightly larger left ventricular end-diastolic diameter and anterior wall thickness compared to *Acomys* (Fig. 1G-J and Supplementary Table 1). We also assessed gross cardiac structure and coronary tree among *Acomys* and *Mus* (C57BL6 and SWR) and found that cardiac structure was not grossly different between the species upon visual comparison of whole hearts along the short- or long-axis (Supplementary Fig. 1G). Using corrosion casting and intra-aortic infusion of Batson’s 17 polymer mixture, we delineated the left coronary anatomy in all three strains. Relevant to permanent coronary ligation, all three animals demonstrated similar course of the left anterior descending artery (LAD) supplying the anterior wall, and of the left circumflex artery supplying the lateral aspect of the left ventricle (Fig. 1K). Coronary blood vessels adopt a hierarchal structure and vascular density leads to enhanced blood supply, angiogenesis and ischemic resistance.^25,26^ Thus, we analyzed the bifurcation points of the coronary tree as it supplies the anterior wall of the left ventricle. These analyses showed a similar density of branching points per major artery across species (Fig. 1L). Finally, we assessed capillary density at the mid left ventricular level between species and observed slightly higher capillary density in *Acomys* and SWR-*Mus* compared to C57BL6-*Mus* (Supplementary Fig. 2). These data demonstrate that cardiac anatomy and physiology are comparable between spiny mice and the two inbred laboratory mouse strains C57BL6 and SWR.

### Adult *Acomys* exhibit a high percentage of cardiomyocytes possessing fetal characteristics

Ventricular cardiomyocyte (CM) phenotype varies across species and chronological age where it can influence the cardiac response to injury.^12,24^ In mammals, the presence of mononuclear diploid CMs are indicative of a ‘young’ heart phenotype that is rarely seen in adult mice.^27, 28^ To examine CM phenotype between species, we isolated ventricular CMs from *Mus* and *Acomys* and examined their size, number of nuclei/CM, nuclear characteristics and ploidy. Similar to a prior report,^29^ *Acomys* CMs were significantly smaller than C57BL6 and were approximately four times more likely to be mononucleated (*Acomys*: 25.9±5.1% vs. C57BL6: 6.3±2.8%, *P* < 0.001; Fig. 2A-C). Similarly, CM nuclei in *Acomys* were smaller in size (Fig. 2D).^30^ We then assessed the ploidy of ventricular CMs across species using 3- dimensional confocal microscopy^31^ and found a significantly higher percentage of diploid CMs in *Acomys* with a corresponding lower percentage of >4N CMs (Fig. 2E). Interestingly, SWR-*Mus* CMs were intermediate between *Acomys* and C57BL6 with smaller size and higher percentage of mononuclear CMs as shown for the inbred strain SWR/J in a previous study.^24^

**Figure 2.**
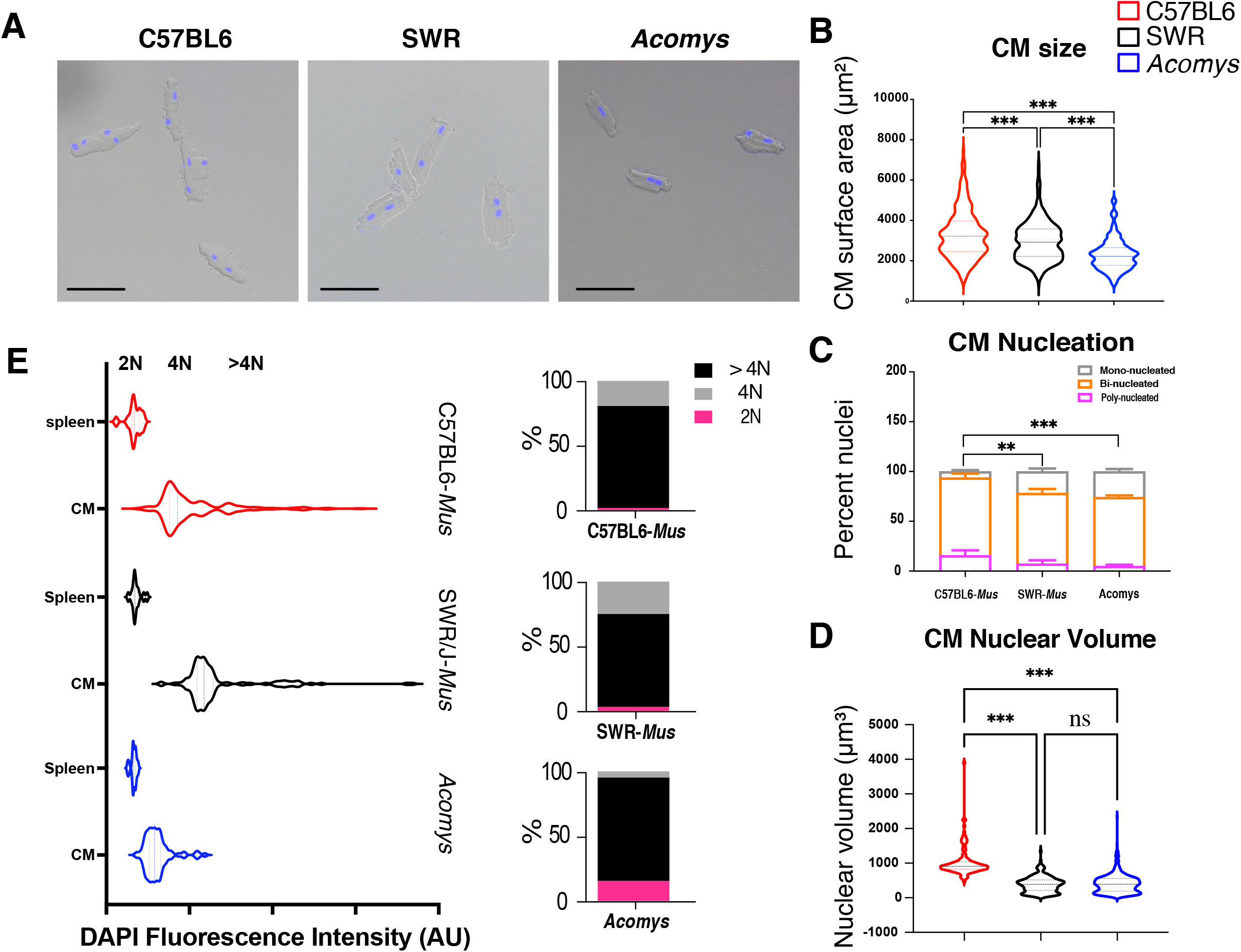
*Acomys* cardiomyocytes exhibit features associated with *Mus* immature cardiomyocytes. (A) Representative images of single-cell ventricular suspension stained with DAPI (blue), identifying small, mononuclear and binucleated cardiomyocytes in *Acomys* and larger binucleated cardiomyocytes in *Mus*-C57BL6 with SWR cardiomyocytes being intermediate in size between C57BL6 and *Acomys* (scale bar = 50 μm) with mononuclear and binucleated cardiomyocytes. (B) Quantitative analyses of cell surface area of *Mus*-C57BL6 and *Acomys* (n = 80 to 120 randomly selected CMs/animal and N = 4 each for *Mus*-C57BL6, *Mus*-SWR and *Acomys*, \*\*\**P* < 0.001 by one-way ANOVA; values are means ± S.E.M). (C) Percentage of mono-, bi- and poly-nuclear cardiomyocytes in C57BL6-, SWR-*Mus* and *Acomys* (n = 80 to 120 randomly selected CMs/animal, N = 4 mice for each group, values are means ± S.E.M, \*\**P*<0.01, \*\*\**P* < 0.001 by One-way ANOVA and Dunnet post-hoc analysis, compared to C57BL6-*Mus*). (D) Nuclear volume among isolated cardiomyocytes showing small nuclear volume in SWR-*Mus* and *Acomys* compared to C57BL6-*Mus* (n = 80 to 120 randomly selected CMs/animal, N = 4 mice per group, values are means ± S.E.M, \*\*\**P* < 0.001 by One-way ANOVA and Dunnet’s post-hoc analysis, compared to C57BL6-*Mus*). (E) Violin plots, on left, demonstrating the distribution of DAPI fluorescence intensity in cardiomyocyte nuclei as assessed using Imaris on 3D (Z-stack) microscopy images. The right panel shows the percentage of CMs within each ploidy category from the total pool of CMs from each species (2N, 4N, and > 4N) (n = 25 to 35 randomly selected CMs/animal, N = 3 mice for each group).

The relatively small cell size and higher mononucleation we observed in *Acomys* CMs suggested a higher percentage of CMs exhibiting immature or fetal characteristics. T-tubule organization is highly regular in mature mouse CMs.^32^ To assess T-tubule organization we stained CMs with di-8-ANEPPS (Fig. 3A) and evaluated transverse and axial density along with T-tubule spacing. *Acomys* showed significantly reduced transverse and long element densities (Fig. 3B and 3C) without a change in resting T-tubule spacing (Fig. 3D). To further explore the physiological implications of these findings, we conducted electrophysiological studies on isolated ventricular CMs from C57BL6-*Mus* and *Acomys*. In mammalian hearts, the expression of low-voltage activated (T-type calcium channels) is an index of the fetal gene program;^33^ therefore, we recorded voltage-dependent I_Ca_ from *Acomys* and *Mus* ventricular myocytes. From a V_hold_ -80mV, a V_test_ step to -25mV elicited little if any discernable current in *Mus* (C57BL6) whereas, at V_test_ -25mV, *Acomys* cardiomyocytes exhibited a transient ‘T-type’ calcium current (I_Ca,T_) (Fig. 3E, upper panel). Longer lasting calcium current (I_Ca,L_) was prominent with larger depolarizations in *Mus* (C57BL6) and *Acomys* (Fig. 3E, lower panel). The current-voltage relationship showed a prominent I_Ca,T_ component with a peak current between V_test_ -20 and -25mV in *Acomys*, but not in *Mus* (C57BL6) (Fig. 3E).

**Figure 3.**
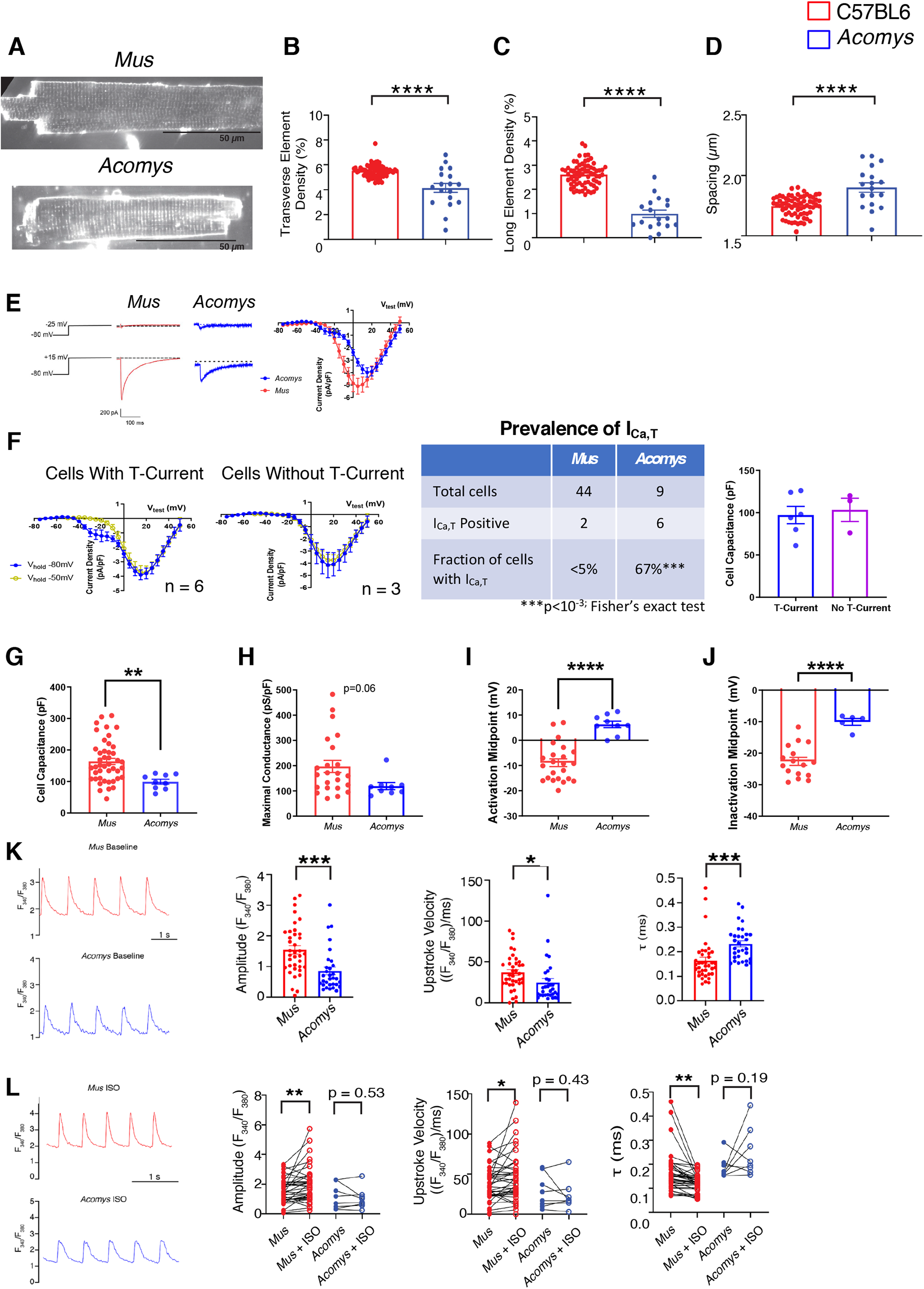
Physiological characteristics of *Acomys* and *Mus* cardiomyocytes. (A) Representative images of isolated cardiomyocytes from C57BL6-*Mus* and *Acomys* stained with di-8-ANEPPS to visualize T-tubules. (B-D) T-tubule analysis shows significantly greater T-tubule density in transverse (B) and axial directions (C) along with wider T-tubule spacing (D) in *Acomys* compared to *Mus* (C57BL6; t=6.2, 12, and 4.4; F=39, 141, and 19, for panels B, C, and D, respectively; *Mus* N=5, n=67; *Acomys* N=3, n=23. (E) Current traces (left) current-voltage relationship (right) elicited by V_test_ -25mV (left, upper) & V_test_ +15mV (left, lower) from V_hold_ -80mV. (F, left). Current-voltage curves from V_hold_ -80mV and -50 mV superimposed for *Acomys* CM exhibiting I_Ca,T_ (A) or no I_Ca,T_. (F, center) I_Ca,T_ expression was heterogeneous but more *Acomys* ventricular CMs showed I_Ca,T_ compared to rare occurrences in *Mus*-C57BL6. (F, right) Cell capacitance was not different for *Acomys* CM with or without I_Ca,T_. (G) Cell capacitance was greater in *Mus*-C57BL6 compared to *Acomys* (t=2.88, F=7.9, \*\**P*<0.01). (H) Maximal conductance density trended greater in *Mus*-C57BL6 compared to *Acomys* (t=2.0, F=7.7, *P*=0.06). (I-J) Voltage-dependent activation and inactivation of I_Ca,L_ was significantly shifted positive for *Acomys* compared to *Mus*-C57BL6 (t=5.8 and 5.2, F=3.7 and 4.5, for panels I and J, respectively, \*\*\*\**P*<0.001). (K) Representative calcium transients from isolated ventricular cardiomyocytes loaded with fura2-AM, *Mus*-C57BL6 (top, red) and *Acomys* (bottom, blue) paced at 1 Hz. Scale bar: 2 seconds. (K, right) Amplitude of the transients (t=3.7, F=1.4, \*\*\**P* = 0.0004). Velocity at which calcium enters the cytosol (upstroke of the transient: t=2.1, F=1.5, \**P*=0.04). (τ) Calcium transient decay (t=3.6, F=1.5, \*\*\**P*=0.0005). (L) Representative calcium transients treated with 100 nM isoproterenol (ISO), *Mus*-C57BL6 (top, red) and *Acomys* (bottom, blue), paced at 1 Hz. Scale bar: 2 seconds. (Left) Before and after ISO Amplitude of the transients (t=2.8 for Mus, P=0.53, t=0.7 for Acomys, \*\**P*= 0.007). (Middle) Before and after ISO Velocity at which calcium enters the cytosol (t=2.5, *P* = 0.02). (Right) Calcium transient decay (t=3.4 for *Mus*, \*\**P* = 0.002; N=7 animals, n=37 cells for C57BL6-*Mus*; and N=4 animals, n=31 cells for *Acomys)*.

To further dissect I_Ca,T_ from I_Ca,L_, we performed current-voltage curve protocols from V_hold_ -50mV to voltage-inactivate the I_Ca,T_ component (Fig. 3F). Current-voltage relationships for I_Ca,T_-expressing *Acomys* cells showed the prominent I_Ca,T_ appearing as a low-voltage activated current for V_hold_ -80mV but not from V_hold_ –50 mV (Fig. 3F, left). 33% of *Acomys* ventricular cardiomyocytes showed no I_Ca,T_ (Fig 3F, center) but there was no observed correlation of cell size to presence of I_Ca,T_ (Fig. 3F, right). Cell capacitance was significantly lower in *Acomys* compared to *Mus* (C57BL6) (Fig. 3G), consistent with our morphometric analysis (Fig. 2B). Additionally, maximal conductance density trended greater in *Mus* (C57BL6) than *Acomys* (Fig. 3H). Voltage-dependent activation and inactivation of I_Ca,L_ was significantly shifted positive for *Acomys* compared to *Mus* (C57BL6) (Fig. 3I-J).

We next explored cytosolic calcium handling. Cytosolic Ca^2+^ transients (CaT) had a larger amplitude (Fig. 3K, left), faster upstroke (Fig. 3K, middle), and more rapid decay (Fig. 3K, right) in *Mus* (C57BL6) compared to that observed in *Acomys*. As with CM morphometrics (Fig. 3A-D) and the prevalent expression of a T-type calcium current (Fig. 3E-F) the smaller, slower CaT segregated with a less mature CM phenotype. Immature ventricular CMs also tend to show reduced β-adrenergic receptor (β-AR) acute responsiveness.^34^ Therefore, we tested the effect of isoproterenol challenge (ISO). To assess acute ISO responsiveness, we compared within cell before – after ISO. *Mus* (C57BL6) showed increased amplitude and more rapid kinetics (Fig. 3L). By contrast, *Acomys* CaT amplitude and upstroke velocity were not significantly different (*P*>0.05). Ca^2+^ re-uptake was accelerated by ISO in *Mus* but not in *Acomys* (Fig. 3L, right). Taken together, these data are consistent with an *Acomys* cardiomyocyte phenotype distinct from that of C57BL6-*Mus*. Moreover, these data demonstrate that a high percentage of *Acomys* CMs exhibit characteristics usually associated with *Mus* fetal CMs.

### *Acomys* hearts demonstrate enhanced cardiac preservation after ischemic injury

*Acomys* can regenerate complex tissue wounds, but the response to myocardial ischemic (MI) injury remains untested. We next sought to compare the cardiac injury response between *Acomys* and the two strains of *Mus* using the permanent LAD-ligation model of myocardial infarction.^35-37^ After injury, all groups showed a comparable drop in cardiac function, reflecting similar injury response as assessed by serial echocardiography (Fig. 4, Supplementary Fig. 3 and Supplementary Table 1). Indeed, there was no significant difference in most of cardiac fucntional parameters at 48 hours after MI (Fig. 4B-F, *P*>0.05). When we assessed the extent of cardiac tissue injury in a subset of animals three days after MI using Mason Trichrome staining, we observed comparable percent infarct between species (Supplementary Fig. 3A). Furthermore, our cardiac MRI imaging showed comparable injury and involvement of the distal anterior wall and apex three days after MI beween species as demonstrated by similar area of late gadolinium enhancement (Supplementary Fig. 4). Interestingly, we observed stabilization in cardiac function and remodeling pararmeters in *Acomys* compared to the progressive deterioration seen in *Mus* strains (C57BL6 and SWR) up to 50 days after injury (Fig. 4B-F). In these analyses, we utilized the delta change from baseline to account for any differences in baseline function between species which allowed us to isolate the analyses to the changes occurring after MI. We observed consistent changes in left ventricular global ejection fraction, as well as measures of left ventricular adverse remodeling such as end-systolic and end-diastolic diameters and volumes (Fig. 4A-F and Supplementary Fig. 3B-F). The enhanced cardiac preservation and reduced adverse cardiac remodeling in *Acomys* was reflected by lower heart/body weight ratio throughout the follow-up period extending to 50 days after MI (Fig. 4G), as well as the aforementioned echocardiographic evidence of smaller ventricular diameters. We also observed significantly lower wet-dry lung weight in *Acomys* compared to *Mus* which is a measure of pulmonary edema and heart failure (Fig. 4H). Importantly, enhanced myocardial preservation in *Acomys* compared to *Mus* was associated with a significant survival advantage (Fig. 4I). As seen in prior reports, the majority of mortality across all species was observed in the first week after MI. However, of those animals that died, while we observed cardiac rupture in 17 C57BL6, we observed none in *Acomys* as visualized in post-mortem necropsies.

**Figure 4.**
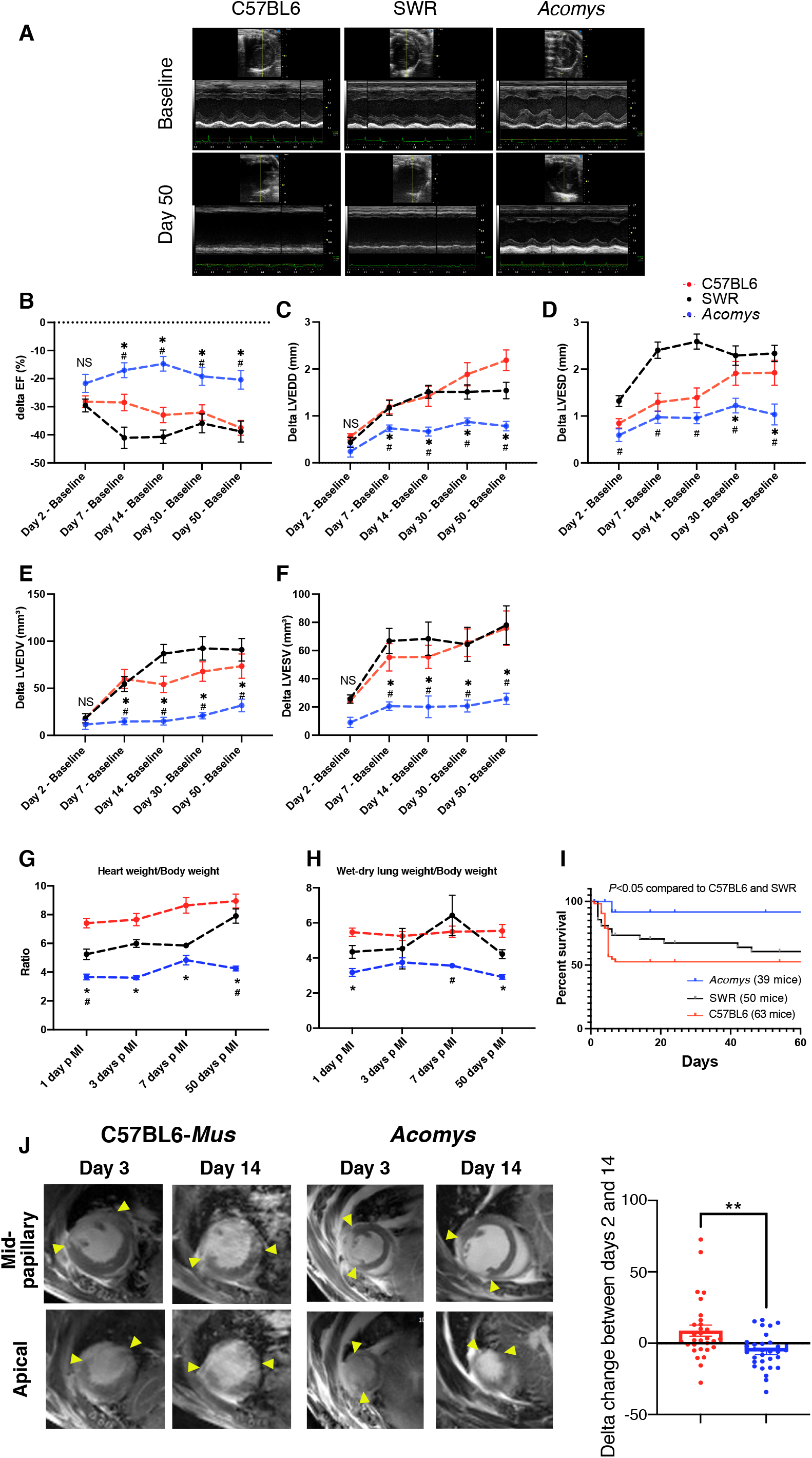
Myocardial ischemic injury response in Acomys and Mus. (A) Representative M-mode echocardiography images of *Mus*-C57BL6) (left), *Mus*-SWR (middle) and *Acomys* (right) at baseline (upper images) and day 50 after MI (lower images). Delta (Δ) ejection fraction (EF) (B), Δleft ventricular end-diastolic diameter (LVEDD) (C), Δleft ventricular end-systolic diameter (LVESD) (D), Δleft ventricular end-diastolic volume (LVEDV) (E), and Δleft ventricular end-systolic volume (LVESV) (F) after injury, calculated as difference between each time point and baseline value in the same animal (N = 16-45 *Mus*-C57BL6, 10-20 *Mus*-SWR and 8-11 *Acomys*) (Values are means ± S.E.M, \**P* < 0.05 by Mixed -effects ANOVA, compared to \**Mus*-C57BL6 or ^#^*Mus*-SWR). (G) Quantitation of heart weight (HW) normalized by body weight (BW) for *Mus*-C57BL6 (N = 2-26 animals), *Mus*-SWR (N = 3-11 animals) and *Acomys* (N = 2-13 animals) at baseline and various timepoints after MI suggesting maintenance of HW/BW ratio in *Acomys* and progressive increase in *Mus* (Values are means ± S.E.M, *P* < 0.05 by two-way ANOVA, compared to \**Mus*-C57BL6 or ^#^*Mus*-SWR). (H) Quantitation of wet-dry lung weight normalized by body weight of *Mus*-C57BL6 (2-26 animals), *Mus*-SWR (3-11 animals) and *Acomys* (1-13 animals) at various timepoints after MI suggesting maintenance of wet-dry lung weight/body weight ratio in *Acomys* and progressive increase in *Mus* (Values are means ± S.E.M, \**P* < 0.05 by two-way ANOVA, compared to \**Mus*-C57BL6 or ^#^*Mus*-SWR). (I) Kaplan-Meier analysis depicting mortality after MI and showing significantly lower mortality in *Acomys* compared with *Mus* (Gehan-Bareslow-Willcoxon test, *P*<0.05). (J) Representative cardiac magnetic resonance (CMR) images showing comparably medium-sized infarct area in both species at three days post MI (arrows show the boundaries of late gadolinium enhancement/injury). The progression of infarct area was significantly slower in *Acomys* compared to *Mus* at 14 days after MI (N= 9-10 animals per group, *t*=3.28, \**P*< 0.05 by paired T-test).

CMR studies demonstrated reduced scar progression between day 3 (D3) and D14 in *Acomys* compared to the continued infarct expansion seen in *Mus* (Fig. 4J). Our long-term follow-up studies reflected significantly smaller infarct size in *Acomys* compared to *Mus* (C57BL6 and SWR) when assessed using Masson’s Trichrome staining 50 days after MI (Fig. 5A). We also saw similar results in our studies quantifying the degree of fibrosis using Picrosirius red staining at 50 days after MI (Supplementary Fig. 3G). Furthermore, the resultant scar in *Acomys* appeared thicker with higher cellular density (Fig. 5C), a finding that could potentially explain the lower rupture and mortality rates. To further explore the fibrous tissue alignment of the thicker scar in *Acomys*, we stained cardiac tissue using Picrosirius Red. While the fiber alignment in the *Mus* strains was typical of a scar with highly parallel and compressed collagen bundles, fiber organization in the center of the *Acomys* scar was more porous with less compression (Fig. 5B). Furthermore, our serial scar analysis at days 7, 17 and 50 after MI demonstrated a unique evolution in infarct size between *Mus* and *Acomys* which corroborated the CMR studies (Fig. 6A-D). In contrast to previous reports,^24^ our studies showed limited recovery of cardiac functional parameters after MI in SWR-*Mus* (Fig. 4B-F). Taken together, our results demonstrate that *Acomys* exhibit ischemic tolerance and enhanced myocardial preservation after ischemic injury compared to the inbred C57BL6- and outbred SWR-*Mus* strains, respectively.

**Figure 5.**
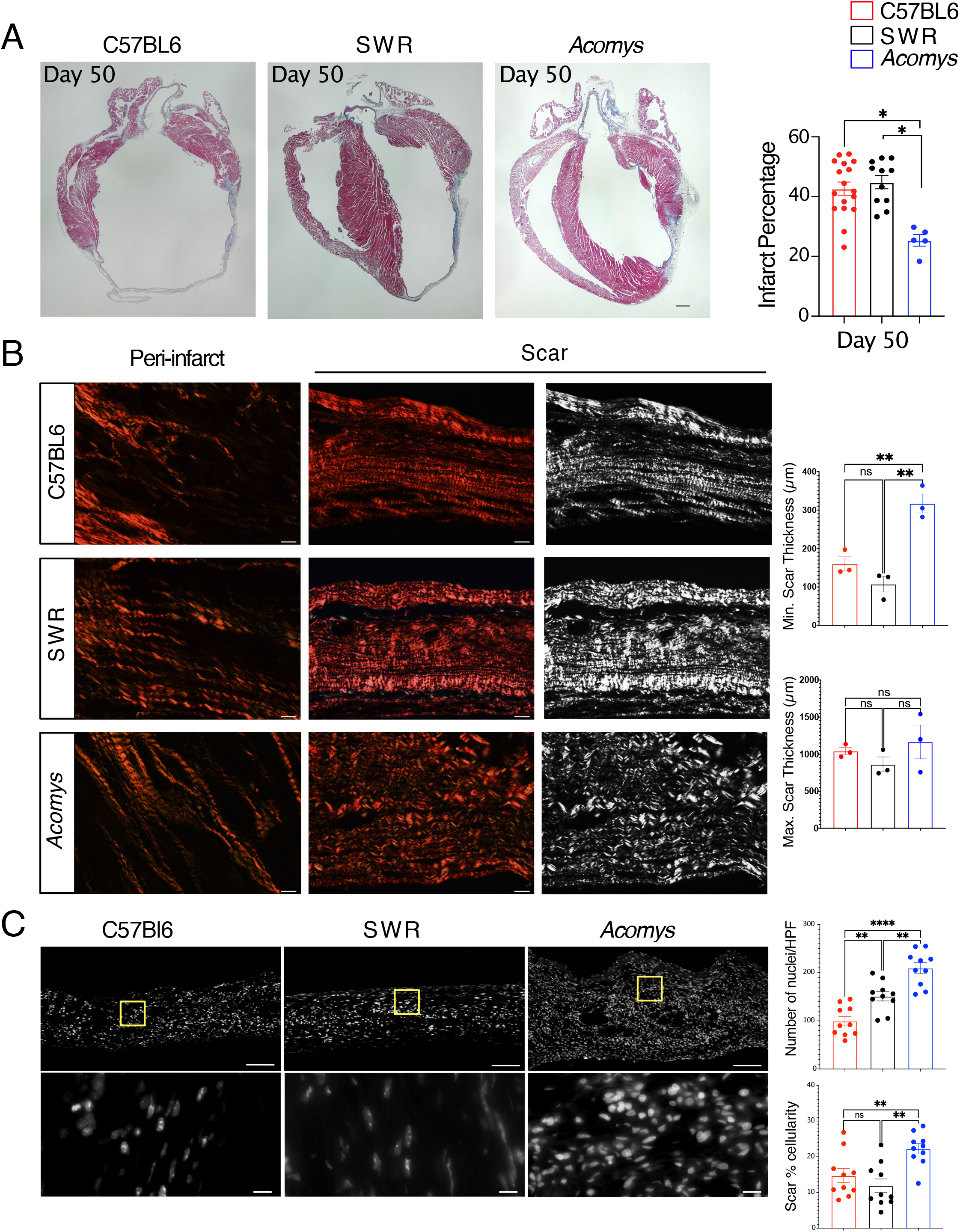
*Acomys* exhibit cardiac tissue preservation in response to myocardial infarction. (A) Representative images of Masson trichrome stained long-axis left ventricular cavity sections of *Mus* (C57BL6 and SWR) and *Acomys* 50 days post-MI (scale bar = 500 μm). Quantification of the infarct size, corresponding to the ratio between infarcted length and left ventricular length showing significantly smaller scar in *Acomys* compared to *Mus* strains (N= 17 *Mus*-C57BL6, 11 *Mus*-SWR, and 5 *Acomys*, values are means ± S.E.M, \**P*< 0.05 by one-way ANOVA and Tukey multiple comparison test). (B) Representative images of picrosirius red stained scar center 50 days after MI across species showing alignment of collagen fibers in *Acomys* compared to *Mus* species. Both *Mus* strains show highly compressed, parallel collagen fibers, while *Acomys* fibers show more porosity between fibers and a wavy organization. Quantification of minimum and maximum scar thickness showing that *Acomys* has on average a comparatively thicker scar compared to *Mus* strains (N= 3 animal/group, values are means ± S.E.M, \*\**P*< 0.01 by one-way ANOVA and Tukey multiple comparison test). There was no statistically significant difference in maximum scar thickness. (C) DAPI images were obtained in the center of the scar and demonstrate higher cellularity in *Acomys* compared to *Mus* strains (scale bar = 100 μm in upper panel and 10 μm in lower panel, yellow boxes indicating insets). Quantitative analysis showed higher density of nuclei and scar percentage cellularity in *Acomys* vs. *Mus* strains (N= 3 animal/group, values are means ± S.E.M, \*\**P*< 0.01 by one-way ANOVA and Tukey multiple comparison test).

**Figure 6.**
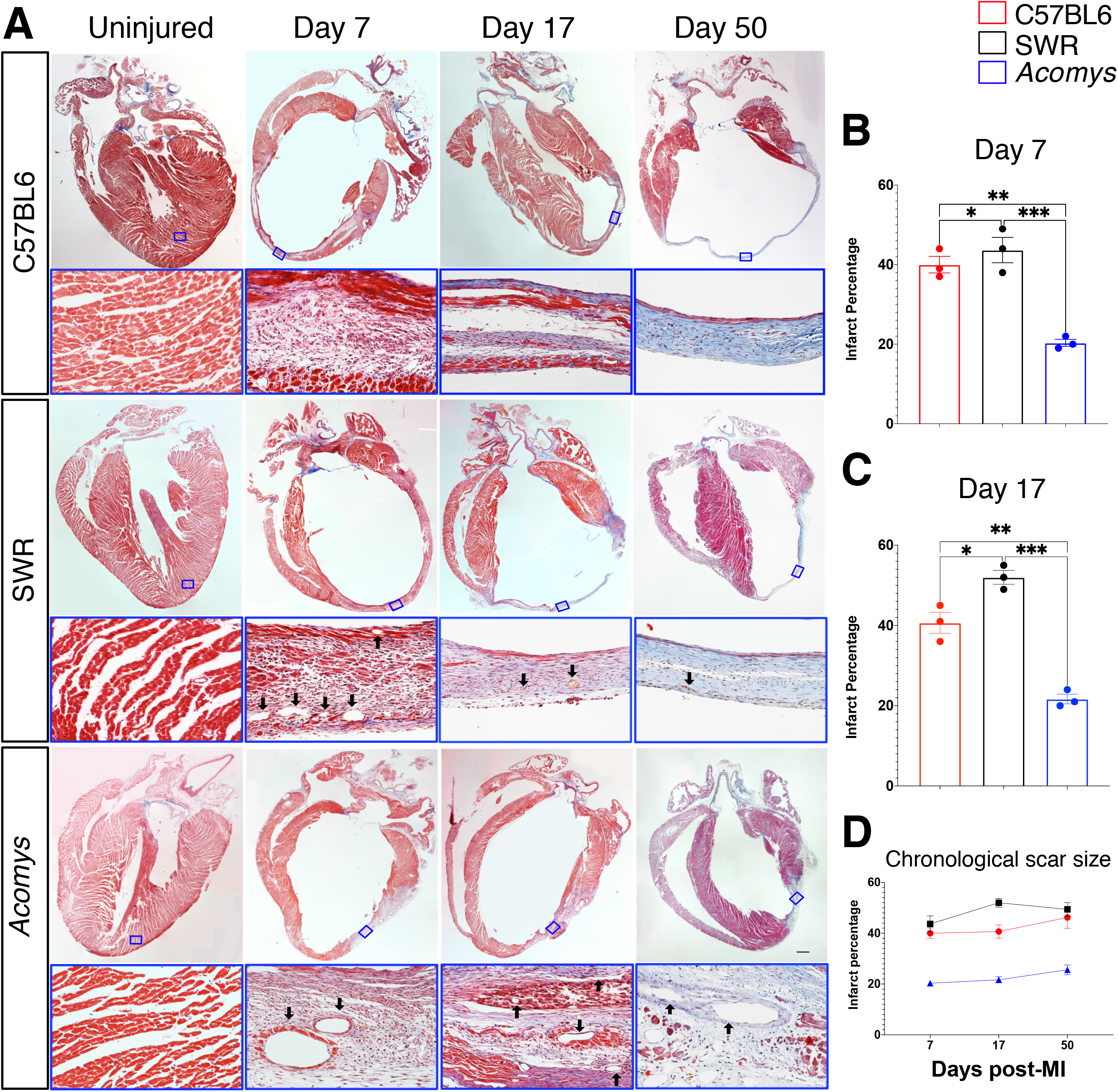
Cardiac scar tissue exhibits limited expansion in *Acomys*. (A) Representative images of Masson trichrome stained long-axis left ventricular cavity sections of *Mus* (C57BL6 and SWR) and *Acomys* in uninjured and MI animals at 7-, 17- and 50-days post-MI showing the progression of scar across species (scale bar = 500 μm). The lower panels represent the boxed areas and show the composition of the scar center at each stage. (B and C) Quantification of the infarct size, corresponding to the ratio between infarcted length and left ventricular length showing significantly smaller scar in *Acomys* compared to *Mus* strains (N= 17 *Mus*-C57BL6, 11 *Mus*-SWR, and 5 *Acomys*, values are means ± S.E.M, \**P*< 0.05 by one-way ANOVA and Tukey multiple comparison test). (D) Delta change in scar size among species showing the reduced delta in *Acomys* compared to *Mus*.

### *Acomys* exhibit rapid and enhanced vascularization of the infarct region after ischemic injury

Blood vessel formation following ischemic injury is essential for functional stabilization. Following ischemic injury, vascular damage ensues in the area surrounding the original infarct region and leads to infarct expansion, development of adverse cardiac remodeling and heart failure.^2,38-40^ Several studies have shown successful reduction of infarct expansion through effective angiogenesis and vascular maturation.^41,42^ We assessed the vascular density in the peri-infarct region in *Acomys* and *Mus* strains and observed an increased density of capillaries (isolectin+ cells) in *Acomys* compared to C57BL6-*Mus* (Fig. 7A). As commonly observed in adult mammals following MI, both *Mus* strains exhibited reduced vascular density in the scar center; by contrast, *Acomys* showed enhanced vascular density extending into the scar center (Fig. 7A). Additionally, enhanced angiogenesis was linked to more proliferating endothelial cells (isolectin+/EdU+) in the peri-infarct region and scar center in *Acomys* compared to both *Mus* strains (Fig. 7B). In support of enhanced angiogenesis, we observed a significantly higher density of mature blood vessels (α-SMA+) in the peri-infarct region in *Acomys* while the presence of these mature vessels was negligible in *Mus* strains (Fig. 7C). Taken together, in response to MI our data support rapid and enhanced angiogenesis followed by vascular maturation in *Acomys* compared to *Mus*. This can explain, at least in part, the ischemic tolerance, reduced infarct expansion and the thicker, cellularly dense scar in *Acomys* compared to *Mus*.

**Figure 7.**
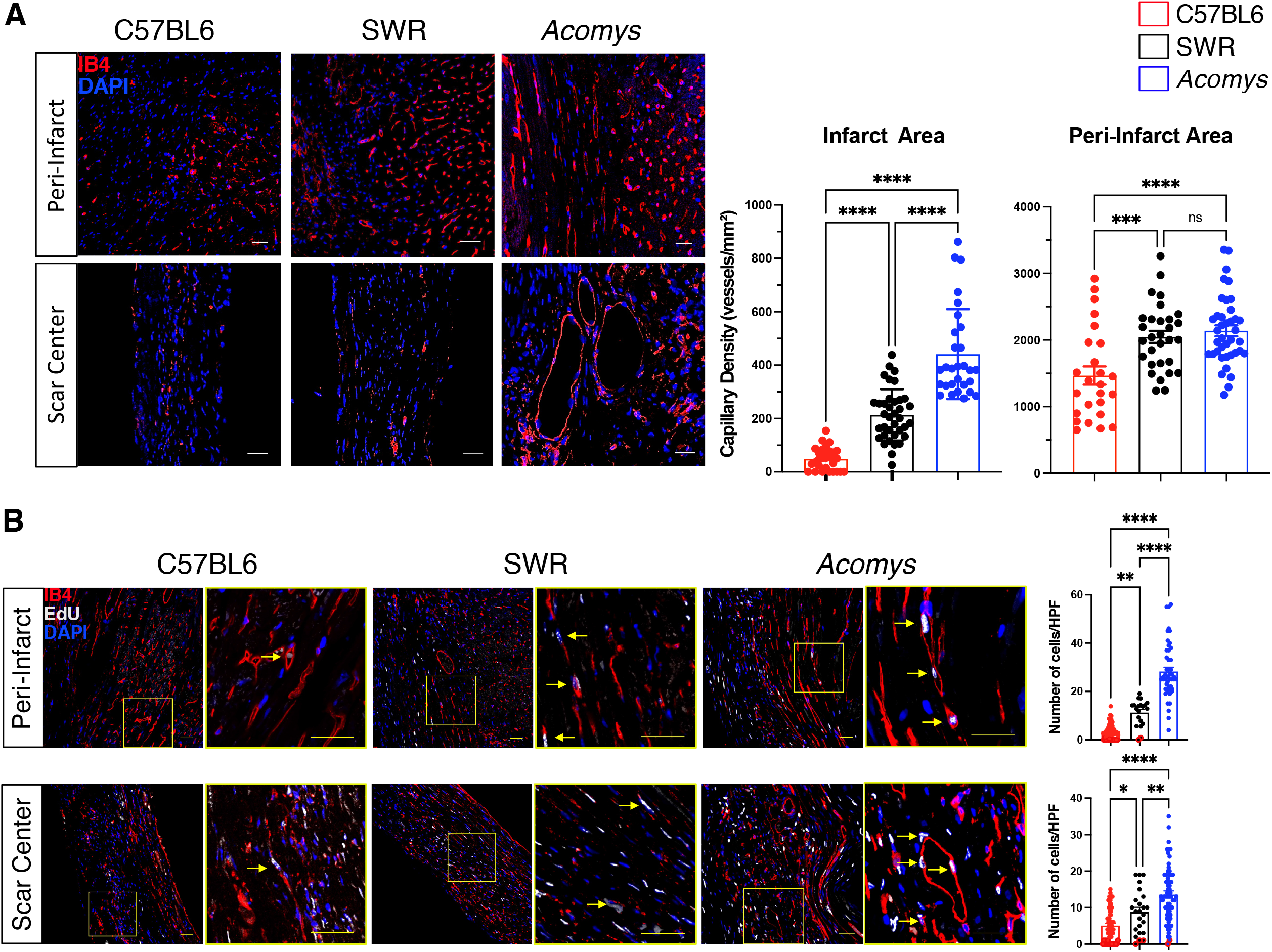
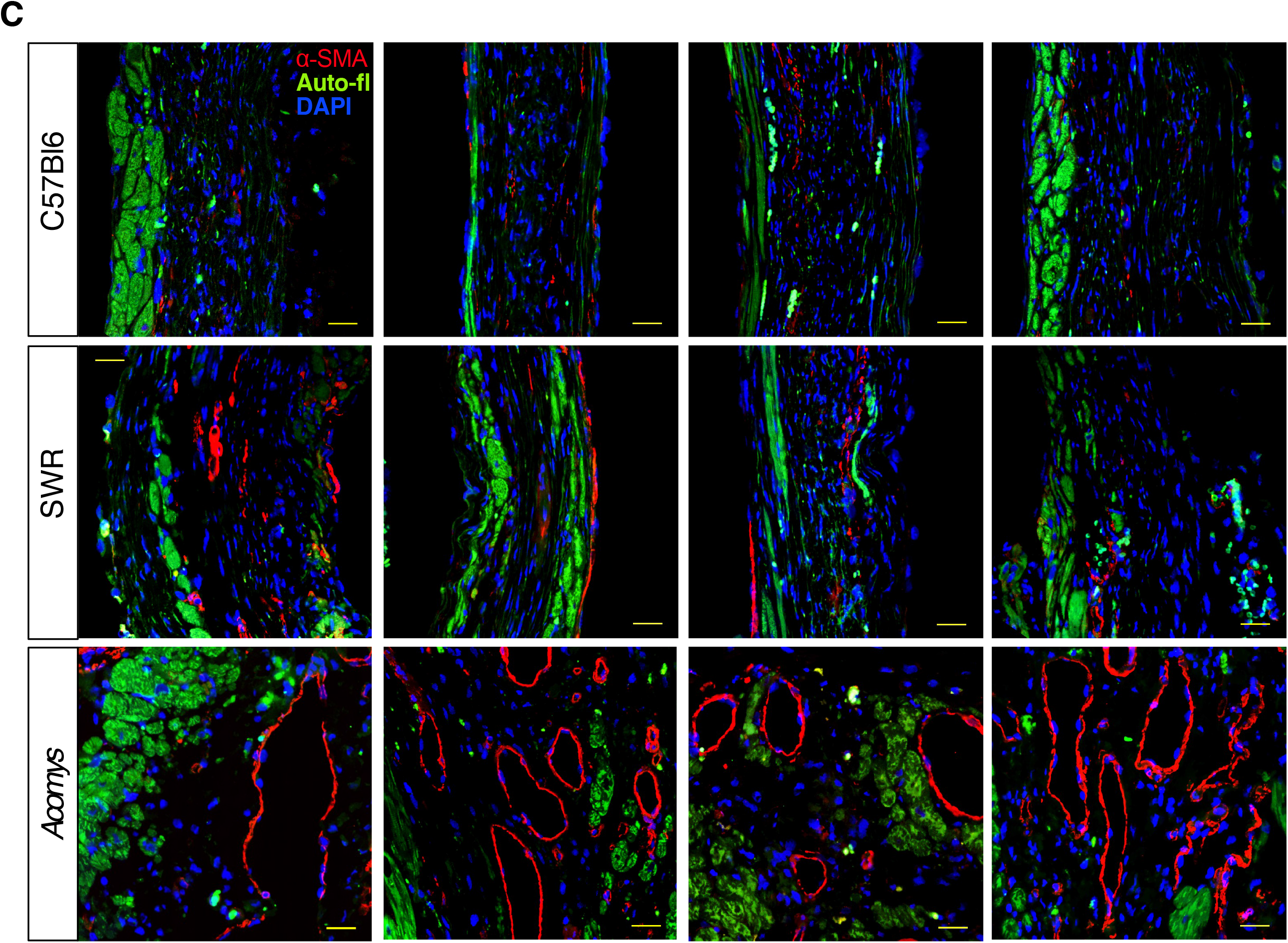
Enhanced angiogenesis and vessel maturation in the infarct region following MI in *Acomys* compared to *Mus* strains. (A) Representative images of isolectin staining (red) defining endothelial cells in the peri-infarct region and the center of the scar at the mid left ventricular cavity level of *Mus* (C57BL6 and SWR) and *Acomys* 50 days post-MI (scale bar = 25 μm). Quantification of capillary density showing significantly higher capillary density in *Acomys* compared to *Mus*-C57BL6 in the peri-infarct region and to both *Mus* strains in the center of the scar (N= 4 *Mus*-C57BL6, 4 *Mus*-SWR, and 5 *Acomys*, values are means ± S.E.M, \**P*< 0.05 by one-way ANOVA and Tukey multiple comparison test). (B) Representative images of isolectin (red) and EdU staining (white) marking proliferating endothelial cells in the peri-infarct region and the center of the scar at the mid left ventricular cavity level of *Mus* (C57BL6 and SWR) and *Acomys* 50 days post-MI (scale bar = 25 μm). Quantification of isolectin+/EdU+ cells showing significantly higher endothelial cell proliferation in *Acomys* compared to the two *Mus* strains in both peri-infarct and scar center areas (N= 4 *Mus*-C57BL6, 4 *Mus*-SWR, and 5 *Acomys*, values are means ± S.E.M, *F* = 28.4, \**P*< 0.05 by one-way ANOVA and Tukey multiple comparison test). (C) Representative images of smooth muscle actin staining (red) in the scar center region showing higher prevalence of medium sized blood vessels in *Acomys* compared to *Mus* strains. Autofluorescence (green) was used to identify the infarct boundary (scale bar = 25 μm).

## Discussion

Spiny mice (*Acomys*) represent a novel mammalian model to explore endogenous cardiac repair consistent with their enhanced regenerative ability for a number of tissues and organs. In this study, we established the anatomical, functional, and cellular characteristics pre- and post-MI in *Acomys* compared to the inbred laboratory mouse strain C57BL6 and the outbred SWR strain. Our results demonstrated comparable cardiac structure, coronary anatomy and functional parameters between the three animals allowing us to appropriately use these animal models for comparative heart injury studies. Interestingly, when comparing cardiomyocytes across species we found that adult *Acomys* possessed a distinct CM phenotype that included features usually observed in immature or fetal *Mus* CMs such as small size, mononucleation and reduced t-tubule density and organization. After ischemic injury, *Acomys* exhibited enhanced ischemic tolerance and significant myocardial preservation, resulting in reduced adverse cardiac remodeling, smaller scar size and, importantly, better survival. Overall, our data supports that *Acomys* show resistance to the type of cardiac damage that compromises other adult mammals, including humans and supports future cardiac studies using this model.

Similar to other highly regenerative vertebrates, *Acomys* has emerged as a bona fide mammalian regeneration model for their ability to naturally regrow complex tissues following full-thickness skin injury^14-19^ and excision of musculoskeletal tissue from the ear pinna.^16,18,43^ In each type of injury, adult *Acomys* were capable of restoring functional tissues, a phenotype that has not been observed in other adult rodents. Our gross anatomical studies established a similar coronary anatomy across species ^44^ and we found that *Acomys* hearts were similar structurally and anatomically to *Mus*. Importantly, *in vivo* cardiac functional parameters were comparable between the three groups. These findings provided the appropriate foundation for the current study and lay the groundwork for follow-up studies that can inform new therapeutic strategies for ischemic heart disease.

Increased mortality and incidence of heart failure following myocardial infarction remains prevalent in modern medicine. While advanced revascularization therapies and mechanical support have slightly improved survival in patients with myocardial infarction, millions develop progressive heart failure every year. Remarkably, *Acomys* exhibited significantly lower mortality after MI compared to both *Mus* strains and this reduction in mortality was related to a reduced incidence of cardiac rupture among *Acomys*. In fact, our histological assessment combined with cardiac MRI studies indicated significant differences in infarct expansion and scar evolution between species. While *Mus* demonstrated progressive infarct expansion, associated with thinning of the anterior wall, *Acomys* showed minimal expansion despite starting with comparable area of injury. These differences in infarct expansion led to favorable cardiac remodeling and functional preservation demonstrated in our echocardiography studies.

This unique phenomenon was associated with a rapid angiogenic response that eventually led to superior ischemic tolerance. *Acomys* scars appeared to be more cellularized and had higher capillary and mature vessel density compared to *Mus*. This could explain the dramatically lower cardiac rupture rate and enhanced survival we observed in *Acomys* including the stabilization of cardiac function in *Acomys* during long-term follow-up. Preclinical and human MI studies have linked enhanced angiogenesis and vessel maturation with lower rates of infarct expansion.^42,45^ Our studies demonstrated superior endothelial cell proliferation and vessel maturation in *Acomys* compared to *Mus* strains which likely accounted for some of the protection seen in our studies.

The mammalian heart loses its reparative capability after the first week of life where heart damage results in permanent loss of myocardium combined with fibrotic scarring and adverse remodeling.^12^ The transition to fibrotic repair is closely linked with the loss of CM proliferation as CMs enter cell cycle arrest and become mononuclear polyploid or multinucleated.^12,46^ However, in other vertebrate species where CMs maintain the lifelong ability to proliferate (e.g., zebrafish and newts), cardiac regeneration is still possible.^46, 47^ One of the hallmarks of regenerative hearts is the relatively high frequency of mononucleated diploid CMs.^46-48^ This evidence suggests that higher proportions of mononuclear diploid CMs are associated with higher regenerative potential following ischemic injury.^24^ Interestingly, our results showed that outbred SWR-*Mus* possessed an intermediate CM phenotype between *Acomys* and C57BL6 with respect to CM size and nuclei number. Despite possessing this CM phenotype, outbred SWR had nearly the same mortality as C57BL6 suggesting that rapid angiogenesis may be a key component to facilitate the protective features conferred by having a larger population of smaller, mononucleate CMs. In fact, cardiac tissue requires vascularization for supporting the high metabolic activity of CMs,^49^ and our current findings support rapid angiogenic invasion into new fibrotic tissue as a mechanism to resist cardiac damage. Future studies are necessary to explore CM behavior across species and how this may relate to enhanced angiogenesis.

There are two types of Ca^2+^ channels in cardiomyocytes, L-type and T-type. L-type Ca^2+^ channels are highly expressed in the adult heart and are important therapeutic targets for the management of various cardiovascular diseases. In contrast, T-type Ca^2+^ channels are rarely found in adults and are present in fetal and early postnatal mice.^50^ In our cell electrophysiology studies, *Acomys* exhibited a higher percentage of T-type Ca^2+^ channels. This feature is consistent with a higher prevalence of phenotypically ‘young’ CMs in *Acomys* compared to *Mus* (C57BL6). Furthermore, immature ventricular cardiomyocytes in *Mus* showed reduced Ca^2+^ transient amplitude, slower kinetics,^51^ and attenuated β-adrenergic receptor responsiveness.^52^ While this unique phenomena in *Acomys* was associated with superior myocardial preservation after ischemic injury, because we did not examine Ca^2+^ channels in outbred SWR mice, we do not know if this feature is associated with the prevalence of small, diploid, mononucleated CMs. However, it is interesting to note that although our CM characterization studies showed that outbred SWR possessed a higher prevalence of small, diploid, mononucleated CMs, this feature alone did not confer the type of enhanced myocardial observed in previous work using inbred SWR/J mice.^24^ This would suggest a synergy between CM phenotype and enhanced angiogenesis with respect to cardiac preservation following MI. Future studies are necessary to examine the proliferative potential of adult CMs in *Acomys* and *Mus* as well as their potential to adequately respond to myocardial injury.

In conclusion, our study presents *Acomys* as a novel model for cardiac research with comparable cardiac structure, function and anatomy to C57BL6- and SWR-*Mus. Acomys* exhibited a distinct cardiomyocyte phenotype and angiogenic response resulting in enhanced recovery and ischemic tolerance after ischemic injury. It is important to note that while we do not see clear evidence of cardiac regeneration, the enhanced myocardial preservation seen in *Acomys* could uncover important therapeutic targets for millions of patients who develop ischemic cardiomyopathy after myocardial infarction. Future studies will focus on the mechanism and potential for these phenotypic differences to foster myocardial preservation and repair instead of scarring and heart failure in models of cardiac disease.

## Acknowledgments

The authors would like to thank Josh Sarli for his help with *Acomys* husbandry and Thomas Wilkop Ph.D at the microscopy core at the University of Kentucky for his assistance with imaging. We thank the Markey Cancer Center and the COBRE histology core for their assistance with the tissue specimen preparation. We thank Binoy Joseph for his assistance in imaging and the Spinal Cord and Brain Injury Research Center for use of their imaging equipment. We thank Beverly Meacham and the University of Kentucky MRISC center for the assistance with cardiac MRI studies.

JS, AWS, AAL, KS, HP, RRD designed the project and experiments. KS, HP, RRD, EG, BMA, BML, DP, HT, and AN performed all the experiments.

KS, HP, RRD, BMA, BML, JS, AAL analyzed the data and KS, HP, JS, AWS, AAL wrote the manuscript. All authors commented on and edited the final version.

## Data Availability

The authors declare that all the data supporting the findings of this study are available within the paper and are contained within supplementary information.

