## Supplementary material for "Adult spiny mice (*Acomys*) exhibit endogenous cardiac recovery in response to myocardial infarction": Suppl material

### Supplementary Information

### Supplementary Methods

#### Animal care

Male *Acomys cahirinus* (6-8 months old, sexually mature animals) were obtained from our in-house breeding colony. Male *Mus musculus* (8-12 weeks, sexually mature animals) C57BL6 were obtained from the Jackson Laboratory, Bar Harbor, ME and outbred SWR were obtained from Charles River. Spiny mice were housed in their own building where they were maintained on an average 12:12 h L:D cycle with exposure to natural light. *Mus* strains were maintained by the University of Kentucky, Division of Laboratory Animal Resources and were exposed to a strict 12:12 h L:D cycle. *Mus* strains were maintained on 18% mouse chow (Tekland Global 2918, Harlan Laboratories, Indianapolis, IN) diet whereas *Acomys* were maintained on 14% mouse chow (Teklad Global 2014, Harlan Laboratories) along with black sunflower seeds. All animal procedures were approved by the University of Kentucky Institutional Animal Care and Use Committee (IACUC #: 2019-3254, 2013-1155 and 2011-0889).

#### Coronary anatomy

Coronary anatomy was visualized using corrosion casting technique. Mice received an i.p. injection of heparin (100 units in 300ul sterile PBS) to prevent coagulation. Ten minutes later, under isoflurane anesthesia, hearts were excised and cannulated with a blunt tip 23g needle and clamp. 10 ml of 1x PBS was perfused intra-aortically, followed by 1.5ml Batson's 17 polymer mixture (2.5ml base monomer + 600ul Catalyst + 3 drops Promoter; Polysciences). Polymer was mixed and ten minutes later was injected over five minutes and then left to cure on ice for 3 hours. Once cured, hearts were briefly washed with ddH2O and then placed in potassium hydroxide (maceration solution) for one hour at room temp. After maceration incubation, hearts were laid with the anterior side up and the left anterior descending artery was imaged using a Nikon Camera (DS-12) after application of several drops of ddH2O to clear overlying tissue. Images were quantified using Imaris® Filament Tracking software. First, the blue channel was extracted and changed to mono. A Gaussian filter was applied, background was subtracted and then semi-automatic tracing was used to identify vessels and branching points. Branching points of all vessels that supply the left ventricular wall were quantified for each species. Five representative images for each species are shown in Figure 1.

**EdU Administration**

To assess angiogenesis, proliferating vessels were quantified in both the peri-infarct and scar regions in mice 17 days post-MI. Mice received a single intraperitoneal injection of 5-ethynyl-2'-deoxyuridine (EdU, 10mg/kg, Cayman Chemical, Ann Arbor, MI) immediately after MI surgery and continuing daily through post-op Day 14. Hearts were collected on Day 17 and processed as indicated in Histology and Immunofluorescence sections. Hearts were stained for Isolectin and EdU with a DAPI counterstain. Images were taken with a 40x oil immersion objective on a Nikon A1 Confocal Microscope in the University of Kentucky Confocal Microscopy facility. 8-10 images per section were taken and only EdU+/Isolectin+ vessels were quantified.

**Tissue collection and Histology**

Tissue (heart and lung) from all species were collected and weighed immediately at sacrifice. The dry lung weight was collected after 3 days of 65° C incubation. Hearts were perfused with PBS (VWR International) followed by 4% PFA (VWR International) fixation via cannulation of the ascending aorta. Hearts were post-fixed overnight at 4°C. Hearts were then sectioned in half long axis at the level of the ligation, and transferred to 70% ethanol until sectioning. Tissues were then paraffin embedded and sectioned. All tissue was sectioned at 5um for immunofluorescence and 8um for Masson’s Trichrome and Picrosirius Red staining.

Masson's Trichrome was imaged using an Olympus BX53 microscope (Olympus, Tokyo, Japan) at 4x and 40x to evaluate scar. The scar size was quantified by examining and calculating the average percent of fibrosis of the total LV area using NIH ImageJ 1.46R software based on Masson's trichrome staining. Picrosirius red staining was performed to visualize scar collagen organization at the peri-infarct and infarct regions. Images of picrosirius stained slides were captured at 40x magnification using Olympus BX53 microscope (Olympus, Tokyo, Japan). The minimum and maximum scar cross sectional length was measured using ImageJ to determine the thickness of scar. DAPI counterstain on ventricular heart sections was performed to analyze cellular density of scar tissue. Images of DAPI stained tissue were taken at 4x and 60x magnification using Olympus BX53 microscope in the University of Kentucky. The number of nuclear in each imaging field was counted using ImageJ. All measurements were analyzed by blinded observers.

**Immunofluorescence**

Immunofluorescence assessments were carried out on deparaffinized and rehydrated sections as previously described. After deparaffinization, slides were washed and underwent TRIS heat induced epitope retrieval for 20 minutes. For EdU/ Isolectin staining, the following EdU staining was performed between the antigen retrieval and protein blocking steps. A reaction cocktail was created using 5ul 2M Tris (pH8.5; Thermo), 2ul 50mM$CuSO_4$ (Fisher), 20ul 0.5M fresh ascorbic acid (Thermo), 2ul Alexa Fluor Azide (0.25mg/ml, Thermo) and 73ul ddH20 per slide. Slides were incubated in the dark in reaction cocktail for 30 minutes at room temperature, then washed in PBS. Protein blocking (3%) was performed for 30 minutes and slides were washed in PBS. Permeabilization was used with 0.3% Triton X-100 for alpha smooth muscle actin staining only. Isolectin stained slides were also blocked for streptavidin/biotin (SP-2002 Vector Labs). After blocking, slides were washed in PBS and incubated with primary antibody: Isolectin B4 (L-2140, Sigma, 1:500) or alpha smooth muscle actin (MAB1420, R&D Systems, 1:50) overnight at 4°C. After washing in PBS, sections were incubated with secondary antibody: streptavidin-Alexa Fluor 568 (S11226, Thermo Fisher,1:500) for Isolectin B4 or Donkey anti mouse Alexa Fluor 568 (ab175700, Abcam, 1:500) for alpha smooth muscle actin for 30 minutes at room temperature. Slides were washed and cover slipped with antifade mounting medium containing DAPI counterstain (H-1200, Vector Labs). 8–15 adjacent areas per section were examined at 40x magnification using Nikon A1 Confocal Microscope in the University of Kentucky Confocal Microscopy facility. Isolectin staining was performed on tissues from day 50 post-MI and was quantified at both the peri-infarct border and the center of the scar and is presented as total capillary density per $mm^2$. Baseline isolectin measurements were taken at a comparable location to the peri-infarct area.

**Cardiomyocyte isolation**

Ventricular cardiomyocytes were isolated as previously described.[24] Briefly, animals received an i.p. injection of heparin (0.3ml of 1000 units/ml) prior to sacrifice. Animals were then anesthetized with 1-3 % Isoflurane. Hearts were excised and immediately perfused on a Langendorff apparatus with a high-potassium Tyrode buffer and then digested with 5 to 7 mg of liberase (Roche Applied Science). After digestion, atria were removed, and ventricular myocytes were mechanically dispersed. Some isolated ventricular cardiomyocytes were used for measuring cell surface area and nuclei, and the others were used for electrophysiological recordings and calcium transients. For electrophysiology and calcium transient measurements control Mus containing $Rad^{fl/fl}$.[24] Some of this control Mus data appeared in reference 19. Calcium concentrations were gradually restored to physiological levels in a stepwise fashion for electrophysiological studies, and only healthy quiescent ventricular myocytes were used for electrophysiological analysis within 12 h.

**Electrophysiological recordings and calcium transients**

$I_{Ca,L}$ was recorded in the whole-cell configuration of the patch clamp technique as previously described.[24] All recordings were performed at room temperature (20 to 22 °C). The pipette solution consisted of (in mmol/liter) 125 Cs-methanosulfonate, 15 TEA-Cl, 1 MgCl2, 10 EGTA, and 5 Hepes, 5 MgATP, 5 phosphocreatine, pH 7.2. Bath solution contained (in mmol/liter) 140 NaCl, 5.4 KCl, 1.2 KH2PO4, 5 Hepes, 5.55 glucose, 1 MgCl2, 1.8 CaCl2, pH 7.4. Once a cell was successfully patched, zero sodium bath solution was introduced into the chamber (mmol/liter) 150 N-methyl-D-glucamine, 2.5 CaCl2, 1 MgCl2, 10 glucose, 10 Hepes, 4-amino-pyridine, pH 7.2. Recordings of isoproterenol response were recorded in zero sodium bath solution containing 300 nM isoproterenol. $I_{Ca,L}$ was recorded from a holding potential ($V_{hold}$) of -50mV. $I_{Ca,T}$ and $I_{Ca,L}$ was recorded from $V_{hold}$ -80mV with 300ms depolarization steps to levels as shown in Figure 3. Activation voltage dependence parameters were obtained by first transforming the peak current voltage relationship to a conductance transform by fitting the ascending phase (typically Vtest +15 to +40mV) to a linear regression to obtain a reversal potential. Using G=I/V the conductance as a function of voltage transform was then fitted to a Boltzmann distribution of the form $G(V)= Gmax/[1+\exp(V_{1/2}/k)]$ where Gmax is the maximal conductance and $V_{1/2}$ is the activation midpoint and $k$ is the slope factor. For steady state availability curve, we pre-pulsed cells to Vpre ranging from -40 to +10 mV in 5 mV increments and recorded a $V_{test}$ to 0 mV. Peak currents were plotted as a function of Vpre, and a Boltzmann distribution was fitted to the resulting curve.

Calcium transients were recorded from ventricular cardiomyocytes loaded with cell permeable Fura-2-AM (Invitrogen). Cardiomyocytes were field stimulated at 1.0 Hz to determine transient amplitude, upstroke velocity, and rate of decay. All measurements were made following >2 minutes of conditioning of 1 Hz-field stimuli to induce steady state. Transients were recorded at 1 Hz. All $Ca^{2+}$ transient / sarcomere dynamic data were analyzed using IonOptix IonWizard 6.3 (IonOptics Inc., Westwood, MA). Background fluorescence (Fbackground) for F380 and F340 were determined from cell-free regions. Data are expressed as F340/380 and were corrected for Fbackground.

**T-tubule quantification**

For T-tubule quantification we used dispersed cardiomyocytes that were not re-exposed to physiological calcium to preserve resting cardiomyocyte length. Cardiomyocytes were incubated in Di-8-ANEPPS (5mM; ThermoFisher, cat#D3167) for 5 minutes, cells were rinsed in low-calcium relaxation buffer, and photographed on a Zeiss5Live laser scanning confocal microscope with a 100x oil immersion objective. Di-8-ANEPPS was excited at 488nm and emission wavelengths >505nm was collected for mid-section slice without visible nuclei. Photomicrographs were analyzed for T-tubule organization using the AutoTT program designed by Drs. Guo and Song.[51] AutoTT allows simultaneous measurement of the longitudinally oriented and transversely oriented T-tubules in each of the ANEPPS-labeled cardiomyocytes.

**Cardiomyocyte surface area and nucleation measurement**

Isolated ventricular cardiomyocytes were fixed in 4% PFA for 10 minutes at 37°C and washed once with PBS by centrifugation at 300 RPM for 3 minutes. Collected cells were stained with DAPI at 1:10,000 dilution for 5 mins at 37°C (H3570, Thermo Fisher Scientific, Waltham, MA) following by another PBS wash prior to mounting on glass slides with ProLong™ Gold Antifade mounting solution (Thermo Fisher, P36934). For surface area quantification, isolated cardiomyocytes were examined using an Olympus IX-71 with DP72 color camera (12.8 megapixel cooled digital color camera). Measurement of CM size was performed using Olympus cellSens software area measuring function to trace the surface of cardiomyocytes. For nucleation analysis, the number of DAPI stained nuclei/CM from different fields were quantified.

### Ploidy and nuclear volume measurement

Cardiomyocyte ploidy analyses were done as previously described with modifications.[30] PFA fixed ventricular cardiomyocytes from each group were used in ploidy and nuclear volume measurement. Confocal z-stacks of randomly selected cardiomyocytes were captured using Nikon Ti, A1 confocal microscope (Nikon, Japan) with a step size of 3 μm. 40x oil immersion objective was utilized for all acquisitions. Ploidy and nuclear volume were measured in z-stacks using Imaris Version 9 software (Bitplane AG, Switzerland). 3D Nuclei were identified and outlined with a standard threshold requirement for all groups using the Imaris cell analysis function to determine nuclear volume and mean DAPI intensity for ploidy analysis. Mean DAPI intensity of spleen cell nuclei from each group were used as reference for diploid (2N) nuclei. Results were plotted as Violin plot using Prism 9. Cardiomyocyte nuclei were determined as diploid if their normalized intensity values were within 1-1.5 times range of reference cells. Nuclei were determined as tetraploid if their intensity values were within 1.5-2.5 times range of reference cell. Polyploidy was assigned if DAPI intensity was greater than 2.5 times range of reference cells.

### Murine model of myocardial infarction

MI surgery was performed as previously described.[23] In brief, we anesthetized mice with 1–3% isoflurane using a small animal vaporizer system. The pain reflex was examined to make sure that the mice were adequately anaesthetized before surgery. A thoracotomy was performed between the 4th and 5th ribs and the pericardial sac was removed. The heart was exposed and pushed out of the chest. The left anterior descending coronary artery (LAD) was identified under direct vision and was permanently ligated 3 mm below its origin using 6–0 silk suture. After LAD ligation, the heart was returned to the intrathoracic space, the Muscles were closed, and the skin was sutured using 4–0 prolen running sutures.

### Echocardiography

A heating pad and rectal temperature probe were utilized to keep body temperature at 37°C during the experiment. Modified parasternal long-axis and short-axis was utilized to determine left ventricular function and volume in M-mode, two-dimensional and Doppler echocardiography modes. We also used M-mode tracings at the mid-papillary level to estimate the systolic and diastolic parameters, and Teichholz formula at end-systole and end-diastole to measure the left ventricle (LV) volumes. All animals were anaesthetized using 1%–3% isoflurane during Echocardiography to maintain a heart rate of 450–500 BPM for all echocardiographic acquisitions.

### Cardiac Magnetic Resonance Imaging

Cardiac magnetic resonance imaging (CMR) was performed on a 7-Tesla ClinScan system (Bruker, Ettlingen, Germany, http://www.bruker.com) equipped with a 4-element phased-array cardiac coil and a gradient system with a maximum strength of 450 mT/m and a maximum slew rate of 4,500 mT/m/s. We acquired whole short-axis stack images from base to apex for comparison with late enhancement images. The short-axis images were planned perpendicular to the four-chamber long-axis image. For late gadolinium-enhanced magnetic resonance imaging, a 0.6 mmol/kg bolus of gadolinium-diethylene triamine pentaacetic acid (Gd-DTPA; Magnevist, Schering Health Care, West Sussex, U.K., http://www.schering.co.uk) was injected using the intraperitoneal route. Imaging was initiated 10 minutes after the injection of Gd-DTPA using an electrocardiographically gated segmented magnetization-prepared fast low-angle shot sequence with a fixed inversion time at 500 ms. The CMR data were analyzed using commercially available postprocessing software: CMR42 (Circle Cardiovascular Imaging, Calgary, Alberta, Canada, http://www.circlecvi.com) and ImageJ (NIH). Infarct length was calculated as percentage of left ventricular length.

### Statistics

Values are expressed as mean ± standard error of mean (SEM). We used student T-test, one-way-ANOVA, repeated measures ANOVA and mixed-effects ANOVA with Tukey corrections to compare data across species as appropriate. The mixed effects model integrates analysis that uses the available data to estimate model parameters. Animals with partial data contribute to the estimation of some model parameters but not others. It is important to note that relying solely on repeated-measures ANOVA will limit the sample size to only animals that survived for the entire duration of the study and could lead to selection bias as we are only selecting a subgroup of animals. Sample sizes, statistical tests and *P* values are indicated in the figures or figure legends. Animal numbers are presented as a range based on the number of animals included in each analysis. These numbers are included in the figure legends. Throughout the analyses, a *P* value $< 0.05$ was considered statistically significant.

### Supplementary Figures

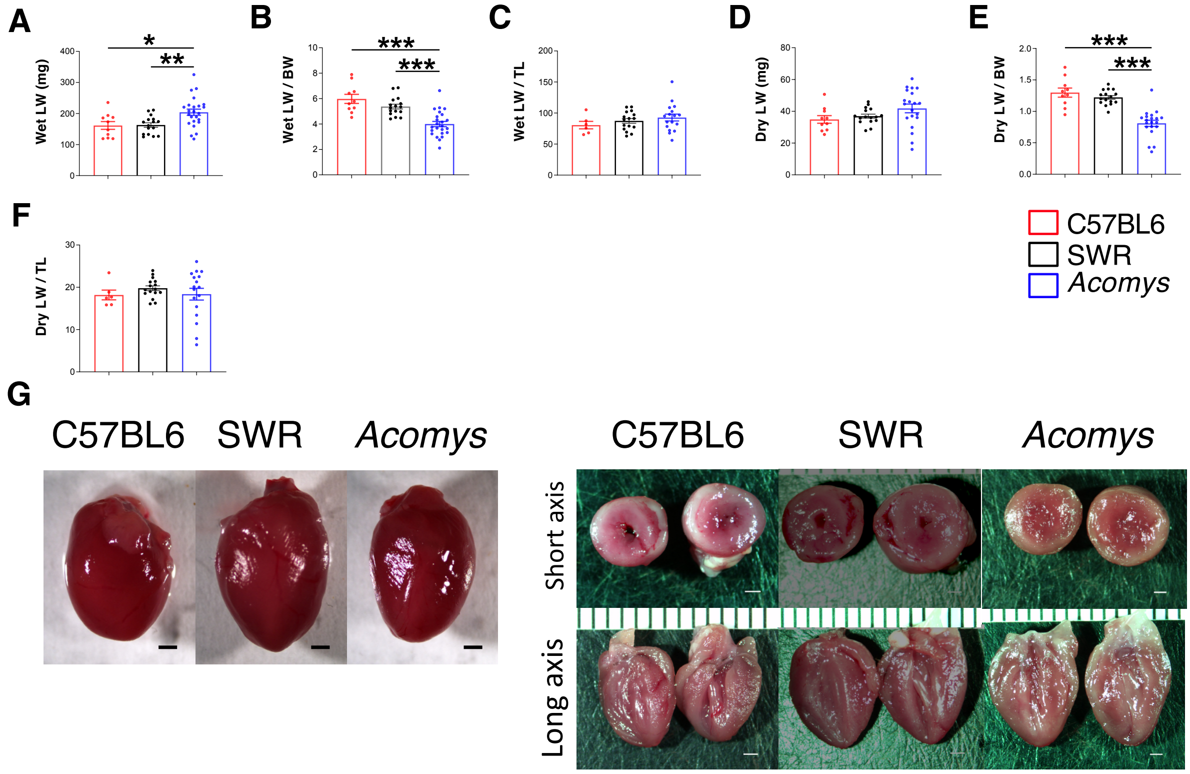


**Supplementary Figure 1. Cardiac function, gravimetric and vascular characterization across mouse strains.** (A-F) Quantitation of dry and wet lung weight of *Mus*-C57BL6 (N = 10), *Mus*-SWR (N = 19) and *Acomys* (N = 22). Analyses demonstrate comparable lung weight across species when normalized by body weight (BW) and tibia length (TL) (values are means ± S.E.M, **P* < 0.05, ***P*<0.01, and ****P*<0.001 by one-way ANOVA and Dunnett correction with Acomys as control). Ejection fraction (F=4.1, **P*<0.05). (G) Representative whole hearts, short axis and long axis *Mus* (C57BL6) (left), *Mus* (SWR) (middle) and *Acomys* (right) at baseline.

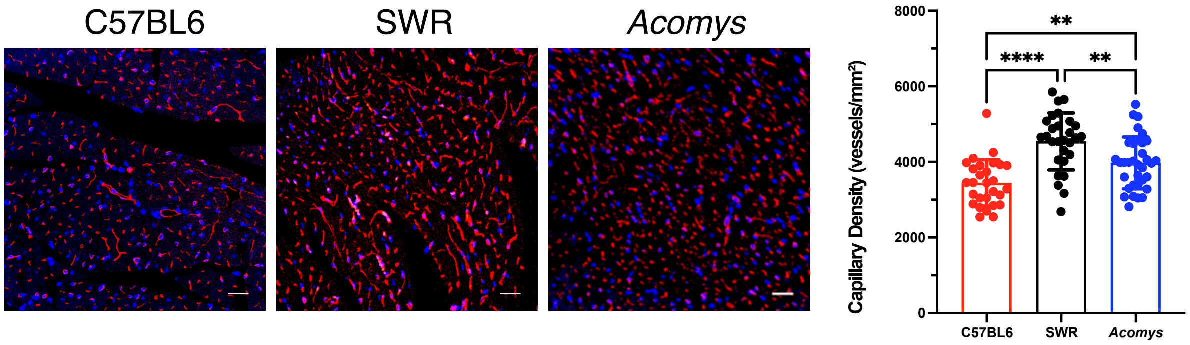


**Supplementary Figure 2. Baseline capillary density across species.** Representative image of isolectin staining of cardiac tissue at baseline in all species. Quantitative analysis showed higher capillary density in SWR-*Mus* compared to C57BL6-*Mus* and *Acomys* (N= 3 animal/group, values are means ± S.E.M, ***P*< 0.01 by one-way ANOVA and Tukey multiple comparison test).

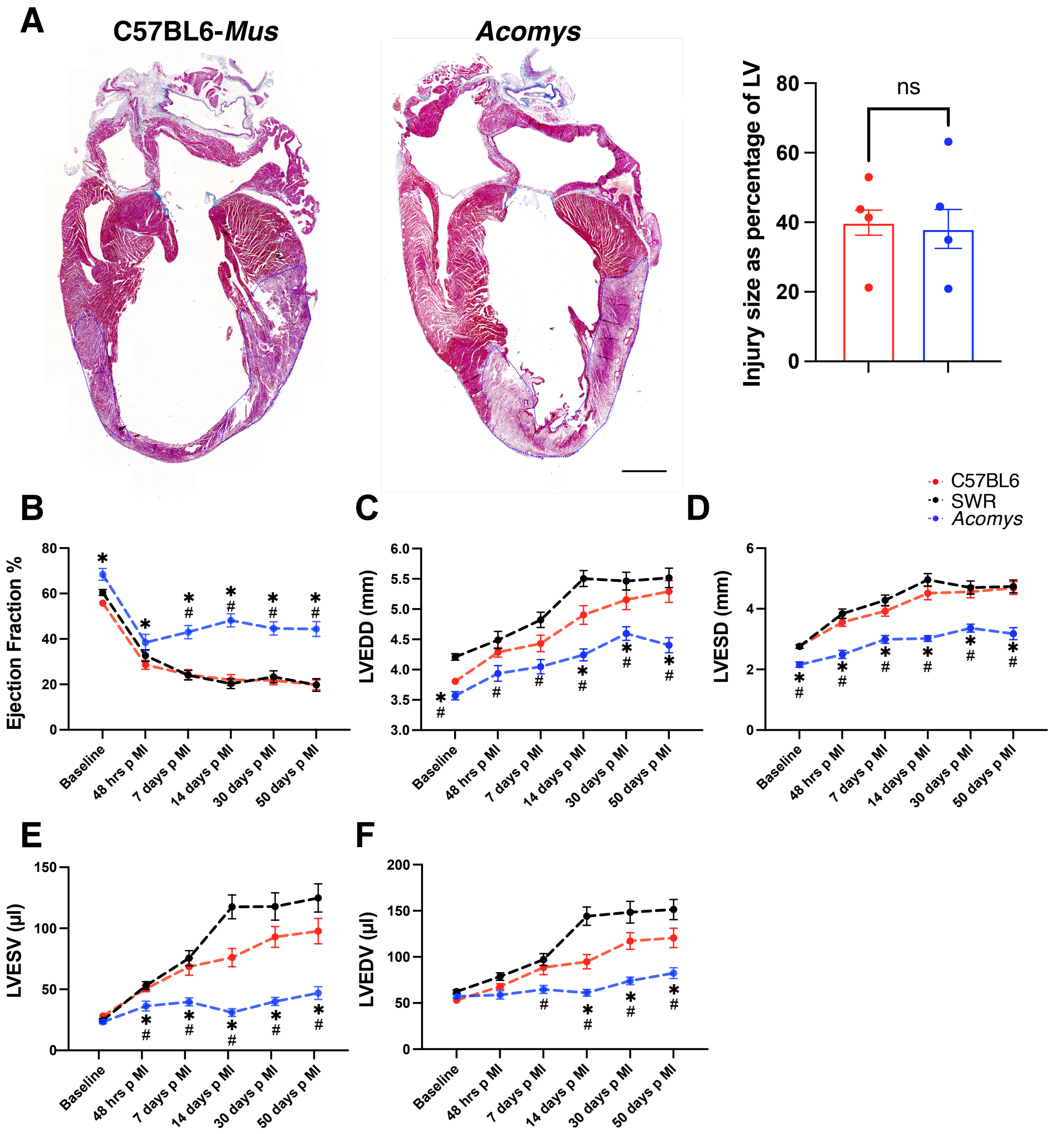


**Supplementary Figure 3. Smaller scar and functional stabilization observed in *Acomys* but not *Mus* after MI.**

Representative images of Masson trichrome stained long-axis left ventricular cavity sections of *Mus* (C57BL6) and *Acomys* at 3 days post-MI (scale bar = 1 mm). Quantification of the size of myocardial injury, showing initial injury in *Acomys* and *Mus* strains (N= 4 C57BL6-*Mus* and 4 *Acomys, P*=0.79 using *t*-test). Echocardiographic recordings of left ventricular ejection fraction (B), left ventricular end-diastolic diameter (LVEDD) (C), left ventricular end-systolic diameter (LVESD) (D), left ventricular end-systolic volume (LVESV) (E), and left ventricular end-diastolic volume (LVEDV) (F) at baseline and 2, 7, 14, 30, 50 days after MI (N = 16-45 *Mus*-C57BL6, 10-20 *Mus*-SWR and 8-11 *Acomys*). These data suggest similar injury pattern early after MI across species with preservation of cardiac function in *Acomys* but not *Mus* strains in long-term functional follow-up (values are means ± S.E.M, $^{*\#}P < 0.05$ by Mixed -effects ANOVA, compared to **Mus*-C57BL6 or $^{\#}$*Mus*-SWR). (G) Representative images of Picrosirius red stained mid left ventricular cavity sections of *Mus* (C57BL6 and SWR) and *Acomys* 50 days post-MI (scale bar = 400 μm). Quantification of fibrosis, corresponding to the ratio between infarcted length and left ventricular length showing significantly smaller fibrosis extent in *Acomys* compared to *Mus* strains (N = 4 *Mus*-C57BL6, 4 *Mus*-SWR, and 5 *Acomys*, values are means ± S.E.M, $F = 45.32$, $^{****}P < 0.0001$ by one-way ANOVA and Tukey multiple comparison test).

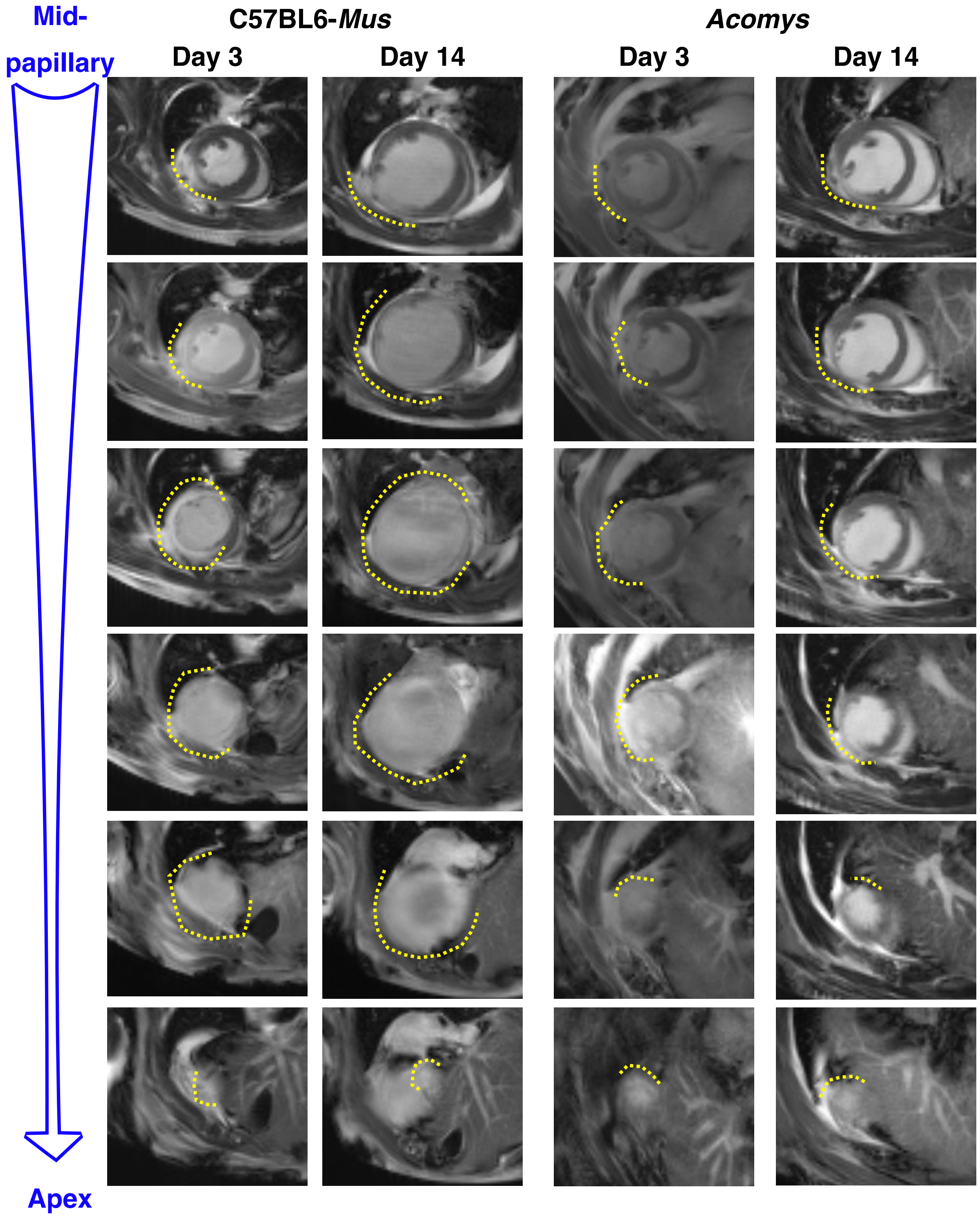


262

263 **Supplementary Figure 4. Comparable early injury and reduced subsequent infarct**
264 **expansion in *Acomys* compared to *Mus*.** Serial representative CMR images from the mid
265 papillary muscle all the way through the distal anterior wall/apex levels in a single C57BL6-
266 *Mus* and *Acomys* showing involvement of the distal anterior wall/apex in both species. While

the scar is stabilized or regressed slightly in Acomys, C57BL6-*Mus* mice show evidence of infarct expansion by 14 days after injury.

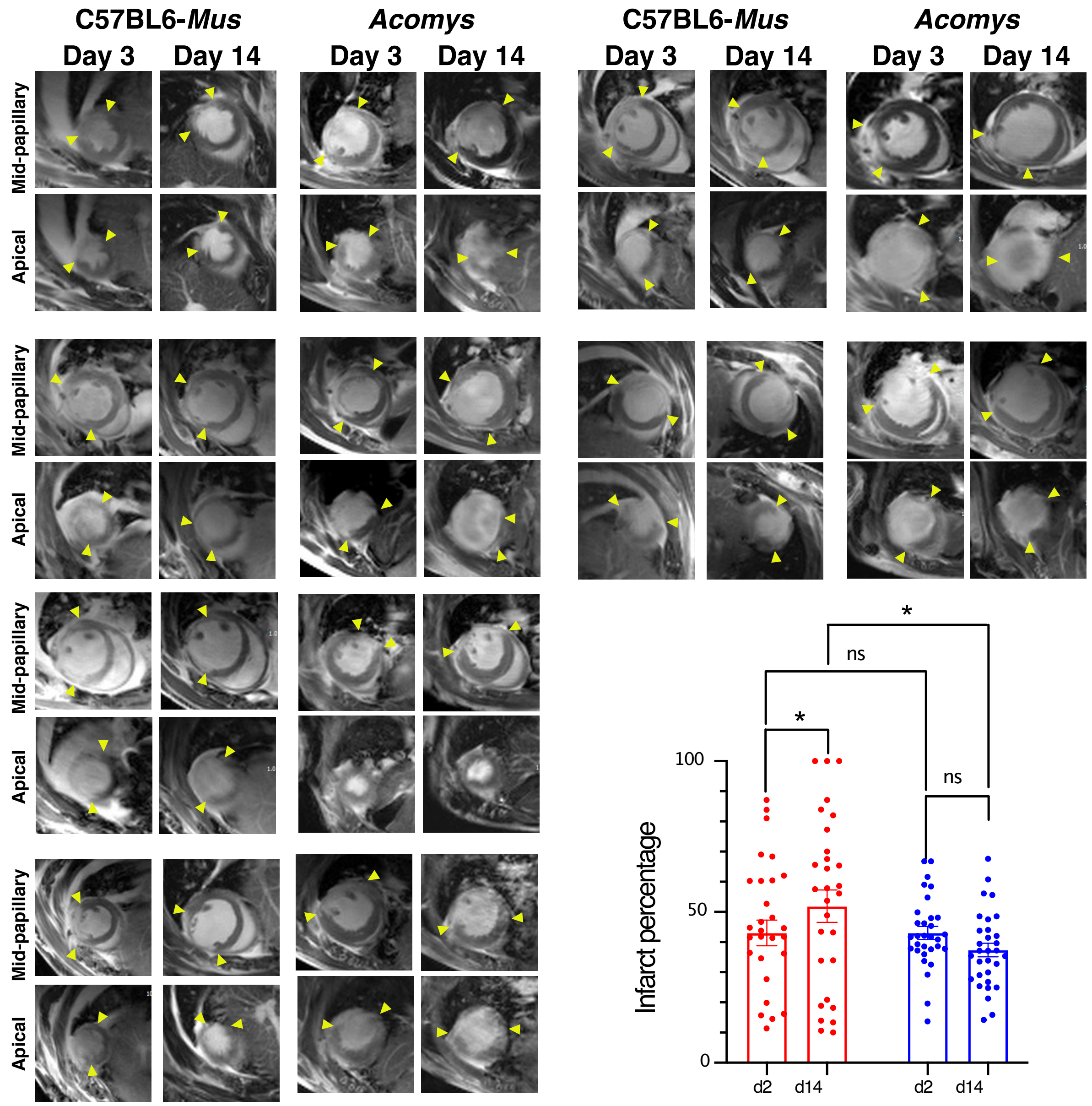


**Supplementary Figure 5**. **Reduced infarct expansion in *Acomys* compared to *Mus*.** (A) Representative cardiac magnetic resonance images showing comparable medium-sized infarct area in both species at 3 days after MI. The progression of infarct area was significantly less in *Acomys* compared to *Mus* at 14 days after MI. The images show transverse sections of the left ventricle starting at the left anterior descending artery ligation site (top) and extending to the apex (bottom). Quantitative assessment of injury size as percentage of left ventricle at 3 and 14 days after MI (N = 9-10 animals per group). All values are means ± S.E.M, *$P < 0.05$ compared to C57BL6-Mus by repeated measures ANOVA.

**Supplemental Table 1: Baseline gross anatomical characteristics and cardiac function pre- and post-myocardial infarction:**

| | | *Mus* (C57BL6) | | | *Mus* (SWR) | | | *Acomys* | | | *P-value* | | |
|---|---|---|---|---|---|---|---|---|---|---|---|---|---|
| | | n | Mean | SEM | n | Mean | SEM | n | Mean | SEM | *Acomys* vs *Mus* (C57BL6) | *Mus* (SWR) vs *Mus* (C57BL6) | *Acomys* vs *Mus* (SWR) |
| **Baseline structural parameters** | | | | | | | | | | | | | |
| BW (mg) | Baseline | 10 | 26.92 | 1.12 | 19 | 40.28 | 0.66 | 22 | 50.26 | 2.18 | <0.0001* | 0.0106* | 0.0019* |
| HW (mg) | Baseline | 10 | 167.58 | 10.41 | 10 | 222.32 | 9.34 | 23 | 231.07 | 12.92 | 0.01* | 0.0143* | >0.9999 |
| HW/BW | Baseline | 10 | 6.22 | 0.29 | 10 | 5.10 | 0.17 | 23 | 4.59 | 0.34 | 0.0045* | 0.1116 | >0.9999 |
| HW/TL | Baseline | 6 | 96.6 | 8.31 | 10 | 117.14 | 5.51 | 17 | 95.81 | 5.59 | >0.9999 | 0.1904 | 0.0191* |
| Wet LW (mg) | Baseline | 10 | 161.4 | 12.55 | 16 | 162.88 | 7.25 | 20 | 204.03 | 9.92 | 0.0435* | >0.9999 | 0.0182* |
| Dry LW (mg) | Baseline | 10 | 34.91 | 2.42 | 16 | 36.96 | 1.26 | 20 | 41.83 | 2.71 | 0.0575 | >0.9999 | 0.2989 |
| Wet LW/BW | Baseline | 10 | 5.98 | 0.36 | 16 | 5.38 | 0.18 | 20 | 4 | 0.2 | 0.0001* | >0.9999 | 0.0004* |
| Dry LW/BW | Baseline | 10 | 1.3 | 0.07 | 16 | 1.22 | 0.03 | 20 | 0.81 | 0.05 | <0.0001* | >0.9999 | <0.0001* |
| Wet LW/TL | Baseline | 6 | 80.62 | 5.99 | 16 | 87.34 | 3.78 | 17 | 92.72 | 5.45 | 0.5355 | >0.9999 | >0.9999 |
| Dry LW/TL | Baseline | 6 | 18.19 | 1.15 | 16 | 19.78 | 0.57 | 17 | 18.38 | 1.39 | >0.9999 | 0.7238 | >0.9999 |
| **Baseline cardiac function and structural characterization** | | | | | | | | | | | | | |
| EF (%) | Baseline | 59 | 55.8 | 1.0 | 35 | 60.5 | 1.3 | 21 | 68.4 | 2.6 | 0.003* | 0.034 | 0.056 |
| FS (%) | Baseline | 24 | 34.10 | 0.77 | 30 | 35.36 | 1.11 | 10 | 35.01 | 1.35 | 0.7274 | 0.3711 | 0.9793 |
| LVEDD (mm) | Baseline | 58 | 3.8 | 0.04 | 35 | 4.3 | 0.05 | 21 | 3.6 | 0.1 | 0.014* | <0.001* | <0.001* |
| LVESD (mm) | Baseline | 60 | 2.8 | 0.04 | 35 | 2.8 | 0.4 | 21 | 2.2 | 0.1 | <0.001* | 0.98 | <0.001* |
| LVAW (mm) | Baseline | 30 | 0.78 | 0.02 | 30 | 0.92 | 0.02 | 10 | 0.88 | 0.02 | 0.044* | <0.0001* | >0.9999 |
| LVPW (mm) | Baseline | 30 | 0.82 | 0.02 | 30 | 0.93 | 0.03 | 10 | 0.90 | 0.04 | 0.1909 | 0.002* | 0.7358 |
| LV mass/BW | Baseline | 30 | 3.31 | 0.07 | 29 | 3.14 | 0.09 | 10 | 2.42 | 0.10 | <0.0001* | 0.6993 | 0.0011* |
| Stroke volume/BW | Baseline | 30 | 1.61 | 0.03 | 29 | 1.01 | 0.04 | 10 | 0.98 | 0.05 | <0.0001* | <0.0001* | >0.9999 |
| **Cardiac recovery after MI** | | | | | | | | | | | | | |
| HW/BW | Day 1 | 3 | 7.402 | 0.3242 | 3 | 5.245 | 0.3627 | 3 | 3.664 | 0.1998 | <0.0001* | 0.0028* | 0.0226* |
| | Day 3 | 5 | 7.653 | 0.4229 | 2 | 5.989 | 0.2677 | 9 | 3.611 | 0.134 | 0.0061* | 0.4014 | 0.1587 |
| | Day 7 | 5 | 8.64 | 0.5366 | 4 | 5.851 | 0.1474 | 5 | 4.839 | 0.3321 | 0.0001* | 0.0167* | 0.4842 |

| | | | | | | | | | | | | | |
|---|---|---|---|---|---|---|---|---|---|---|---|---|---|
| | Day 50 | 26 | 8.947 | 0.4823 | 12 | 7.9 | 0.4953 | 7 | 4.255 | 0.1658 | <0.0001* | 0.1423 | <0.0001* |
| Wet-Dry LW/BW | Day 1 | 8 | 5.469 | 0.2311 | 3 | 4.349 | 0.363 | 7 | 3.178 | 0.2201 | 0.004* | 0.43 | 0.412 |
| | Day 3 | 6 | 5.253 | 0.2493 | 5 | 4.53 | 1.155 | 6 | 3.752 | 0.2609 | 0.123 | 0.643 | 0.6 |
| | Day 7 | 5 | 5.449 | 03208. | 3 | 6.425 | 1.152 | 5 | 3.561 | 0.1435 | 0.06 | 0.608 | 0.011* |
| | Day 50 | 26 | 5.546 | 0.3638 | 20 | 4.222 | 0.2496 | 7 | 2.917 | 0.1288 | <0.0001* | 0.003* | 0.071 |
| Infarct length (%) | Day 50 | 17 | 42.67 | 2.21 | 11 | 44.85 | 2.22 | 5 | 25.37 | 1.94 | 0.0006* | 0.7666 | 0.0003* |
| **Cardiac function after MI** | | | | | | | | | | | | | |
| EF (%) | Baseline | 59 | 55.8 | 1.0 | 35 | 60.5 | 1.3 | 21 | 68.4 | 2.6 | 0.003* | 0.034* | 0.056* |
| | Day 2 | 34 | 28.6 | 1.9 | 14 | 32.7 | 2.6 | 11 | 38.4 | 3.6 | 0.048* | 0.99 | 0.528 |
| | Day 7 | 25 | 24.3 | 2.2 | 25 | 24.03 | 2.1 | 18 | 43.01 | 2.9 | 0.013* | 0.99 | 0.53 |
| | Day 14 | 32 | 21.95 | 2.3 | 20 | 20.3 | 2.2 | 19 | 48.2 | 2.9 | <0.0001* | 0.99 | <0.0001* |
| | Day 30 | 31 | 21.8 | 1.9 | 19 | 23.3 | 2.6 | 22 | 44.6 | 2.9 | <0.0001* | 0.99 | 0.002* |
| | Day 50 | 31 | 19.8 | 2.1 | 17 | 19.7 | 2.8 | 19 | 44.4 | 3.3 | <0.001* | 0.6933 | 0.002* |
| LVEDD (mm) | Baseline | 58 | 3.8 | 0.04 | 35 | 4.3 | 0.05 | 21 | 3.6 | 0.1 | 0.014* | <0.001* | <0.001* |
| | Day 2 | 34 | 4.3 | 0.08 | 14 | 4.5 | 0.2 | 11 | 3.9 | 0.1 | 0.08 | 0.44 | 0.02* |
| | Day 7 | 24 | 4.4 | 0.1 | 25 | 4.8 | 0.1 | 21 | 4.1 | 0.1 | 0.09 | 0.1 | 0.03* |
| | Day 14 | 29 | 4.9 | 0.2 | 20 | 5.5 | 0.1 | 21 | 4.2 | 0.1 | 0.002* | 0.01 | <0.001* |
| | Day 30 | 29 | 5.2 | 0.2 | 19 | 5.5 | 0.2 | 22 | 4.6 | 0.1 | 0.02* | 0.34 | <0.001* |
| | Day 50 | 29 | 5.3 | 0.2 | 17 | 5.5 | 0.2 | 19 | 4.4 | 0.1 | <0.001* | 0.62 | <0.001* |
| LVESD (mm) | Baseline | 60 | 2.8 | 0.04 | 35 | 2.8 | 0.4 | 21 | 2.2 | 0.1 | <0.001* | 0.98 | <0.001* |
| | Day 2 | 34 | 3.6 | 0.1 | 14 | 3.8 | 0.6 | 11 | 2.5 | 0.1 | <0.001* | 0.36 | <0.001* |
| | Day 7 | 26 | 3.9 | 0.2 | 25 | 4.3 | 0.9 | 20 | 2.99 | 0.1 | <0.001* | 0.29 | <0.001* |
| | Day 14 | 30 | 4.5 | 0.2 | 20 | 4.95 | 0.9 | 21 | 3.02 | 0.09 | <0.001* | 0.31 | <0.001* |
| | Day 30 | 30 | 4.6 | 0.2 | 19 | 4.7 | 0.95 | 22 | 3.4 | 0.1 | <0.001* | 0.89 | <0.001* |
| | Day 50 | 30 | 4.7 | 0.2 | 17 | 4.73 | 0.9 | 18 | 3.2 | 0.2 | <0.001* | 0.98 | <0.001* |
| LVESV (μL) | Baseline | 48 | 28.2 | 1.3 | 35 | 24.7 | 1.1 | 35 | 23.3 | 1.6 | 0.054 | 0.12 | 0.77 |
| | Day 2 | 34 | 50.5 | 2.5 | 14 | 53.01 | 3.3 | 17 | 36.3 | 4.01 | 0.015* | 0.8 | 0.009* |
| | Day 7 | 34 | 68.6 | 6.9 | 25 | 75.5 | 6.2 | 24 | 39.8 | 3.3 | 0.001* | 0.74 | <0.001* |
| | Day 14 | 33 | 76.01 | 7.5 | 20 | 117.6 | 9.8 | 20 | 31.03 | 3.2 | <0.001* | 0.004* | <0.001* |
| | Day 30 | 33 | 92.9 | 8.5 | 19 | 117.9 | 11.2 | 22 | 40.1 | 3.3 | <0.001* | 0.19 | <0.001* |

| | | | | | | | | | | | | | |
|---|---|---|---|---|---|---|---|---|---|---|---|---|---|
| | Day 50 | 33 | 97.7 | 10.4 | 17 | 124.8 | 11.6 | 19 | 46.97 | 5.3 | <0.001* | 0.20 | <0.001* |
| LVEDV (μL) | Baseline | 45 | 53.01 | 1.9 | 35 | 62.4 | 2.02 | 35 | 57.2 | 2.02 | 0.29 | 0.004* | 0.18 |
| | Day 2 | 34 | 67.99 | 2.9 | 14 | 78.7 | 4.03 | 17 | 58.8 | 4.5 | 0.22 | 0.09 | 0.008* |
| | Day 7 | 32 | 88.5 | 7.71 | 25 | 97.1 | 6.42 | 24 | 64.8 | 4.4 | 0.03* | 0.67 | 0.004* |
| | Day 14 | 31 | 94.8 | 7.8 | 20 | 144.1 | 9.9 | 20 | 61.1 | 3.7 | 0.001* | 0.001* | <0.001* |
| | Day 30 | 31 | 117.2 | 8.95 | 19 | 148.5 | 11.8 | 22 | 74.03 | 4.1 | <0.001* | 0.1 | <0.001* |
| | Day 50 | 31 | 120.6 | 10.5 | 17 | 151.3 | 10.9 | 19 | 82.4 | 5.8 | 0.007* | 0.12 | <0.001* |

BW; body weight, HW; heart weight, TL; tibial length, LW; lung wight, EF; ejection fraction, LVEDD; left ventricular end-diastolic diameter, LVESD; left ventricular end-systolic diameter, LVAW; left ventricular anterior wall, LV mass; left ventricular mass (*P < 0.05).
